# Genomes of trombidid mites reveal novel predicted allergens and laterally-transferred genes associated with secondary metabolism

**DOI:** 10.1101/259044

**Authors:** Xiaofeng Dong, Kittipong Chaisiri, Dong Xia, Stuart D. Armstrong, Yongxiang Fang, Martin J. Donnelly, Tatsuhiko Kadowaki, John W. McGarry, Alistair C. Darby, Benjamin L. Makepeace

**Author notes:** Contributed equally (Alistair C. Darby, Benjamin L. Makepeace). Correspondence address: Ben Makepeace, Department of Infection Biology, Institute of infection & Global Health, Liverpool Science Park IC2, 146 Brownlow Hill, Liverpool L3 5RF, United Kingdom.

## Abstract

**Background:** Trombidid mites have a unique lifecycle in which only the larval stage is ectoparasitic. In the superfamily Trombiculoidea (“chiggers”), the larvae feed preferentially on vertebrates, including humans. Species in the genus *Leptotrombidium* are vectors of a potentially fatal bacterial infection, scrub typhus, which affects 1 million people annually. Moreover, chiggers can cause pruritic dermatitis (trombiculiasis) in humans and domesticated animals. In the Trombidioidea (velvet mites), the larvae feed on other arthropods and are potential biological control agents for agricultural pests. Here, we present the first trombidid mites genomes, obtained both for a chigger, *Leptotrombidium deliense*, and for a velvet mite, *Dinothrombium tinctorium*.

**Results:** Sequencing was performed using Illumina technology. A 180 Mb draft assembly for *D. tinctorium* was generated from two paired-end and one mate-pair library using a single adult specimen. For *L. deliense*, a lower-coverage draft assembly (117 Mb) was obtained using pooled, engorged larvae with a single paired-end library. Remarkably, both genomes exhibited evidence of ancient lateral gene transfer from soil-derived bacteria or fungi. The transferred genes confer functions that are rare in animals, including terpene and carotenoid synthesis. Thirty-seven allergenic protein families were predicted in the *L. deliense* genome, of which nine were unique. Preliminary proteomic analyses identified several of these putative allergens in larvae.

**Conclusions:** Trombidid mite genomes appear to be more dynamic than those of other acariform mites. A priority for future research is to determine the biological function of terpene synthesis in this taxon and its potential for exploitation in disease control.

## Background

The Acari (mites and ticks) is the most speciose group within the subphylum Chelicerata, with approximately 55,000 described species in both terrestrial and aquatic habitats, and an estimated total diversity of up to 1 million species [1]. This assemblage is paraphyletic and is composed of two major divisions, the Parasitiformes and the Acariformes, which both contain species of medical, veterinary and agricultural importance. For example, the Parasitiformes harbours predatory mites used in the control of agricultural pests (*e.g., Metaseiulus occidentalis*); ectoparasites of honey bees (*Varroa destructor* and *Tropilaelaps mercedesae*) that transmit pathogenic viruses; and most famously, the ticks (Ixodida). The Acariformes includes major ectoparasites and sources of allergens for humans and other animals, such as the scabies mite (*Sarcoptes scabiei*) and the dust mites (*Dermatophagoides* spp. and *Euroglyphus maynei*).

The role of lateral gene transfer (LGT) in the evolution of animals has remained controversial since the first metazoan genomes were sequenced. Despite the explosion of new genomic resources across a wider variety of metazoan phyla in recent years, some claims of large-scale LGT in animal genomes have been shown to be the result of flawed data-analysis methods that failed to exclude sequences from bacterial contaminants [2-5]. Attempts to conduct meta-analyses across diverse metazoan genomes using consistent criteria have also been criticized for reliance on unsound assumptions [6, 7]. However, it is irrefutable that several metazoan taxa are reliant on functions obtained via LGT for essential physiological processes, including digestion of complex and/or toxic materials (especially in the case of herbivores and plant parasites) and the avoidance of host defences [8-11]. One group of acariform mites, the spider mites (superfamily Tetranychoidea: order Trombidiformes; Fig. 1), are major pests of various crops and rely on well-characterised lateral gene transfers of β-cyanoalanine synthase (from bacteria) and carotenoid biosynthesis enzymes (from fungi) to detoxify hydrogen cyanide in their diet [12] and to control diapause [13], respectively. Currently, it is unknown whether LGT is a key feature of the evolution of the spider mites only, or is more widespread among the Trombidiformes.

**Figure 1:**
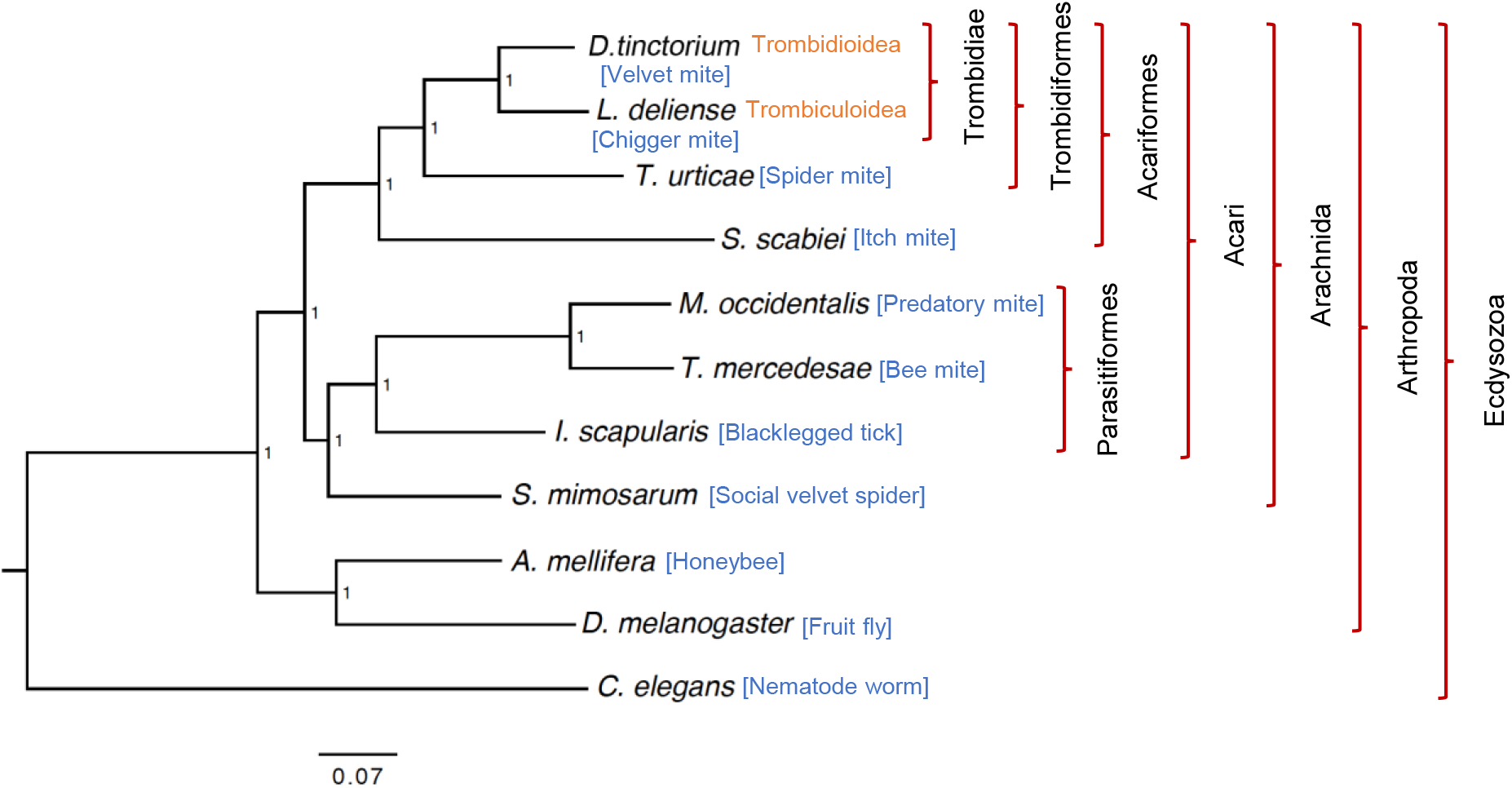
Phylogenetic tree based on the amino acid sequences of 527 one-to-one orthologous genes in 11 species of Ecdysozoa using Bayesian methods. The taxonomy of the trombidid mites follows the scheme of Lindquist *et al.* [14].

In addition to the Tetranychoidea, the Trombidiformes contains two other superfamilies of economic or clinical importance, the Trombidioidea (the velvet mites) and Trombiculoidea (the “chiggers” and related groups) [14]. These two taxa, known collectively as trombidid mites (Fig. 1), have a unique natural history among arthropods in that only the larval stage is ectoparasitic, whereas the nymphs and adults are predators of other arthropods (Fig. 2). However, the Trombidioidea and Trombiculoidea differ in their host preferences. Larvae of the Trombidioidea are exclusively parasites of other arthropods and some species feed on insects of medical, veterinary and agricultural importance, including mosquitoes [15], the New World screwworm fly [16], and aphids [17] (Fig. 2). On dipteran hosts, heavy infestations can reduce flight ability, while certain aphid species can be killed in a few days by as few as two feeding larvae [18, 19].

**Figure 2:**
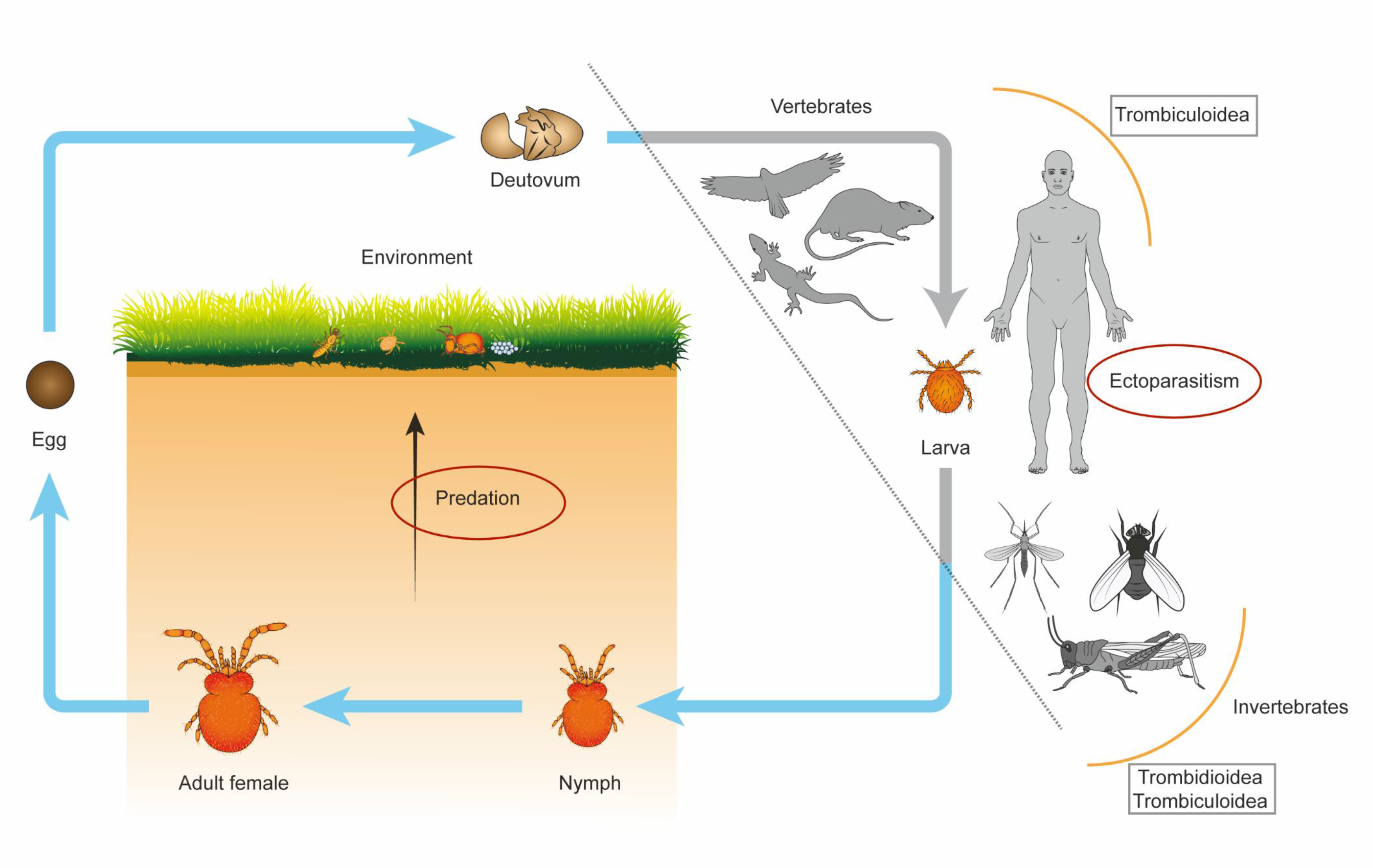
Simplified lifecycle of trombidid mites. In the Trombidioidea (velvet mites), the larvae are parasitic on other arthropods; whereas in the Trombiculoidea (“chiggers”), the larvae feed on a variety of vertebrates or (more rarely) other invertebrates. The deutonymph and adult are free-living, edaphic stages that predate soft-bodied arthropods (*e.g.*, termites, springtails and other mites) or consume their eggs. Trombidid eggs are laid in the environment and produce questing larvae that congregate and seek a host. For clarity, only the deutovum and active instars are shown; the protonymph (between the larva and deutonymph stages) and the tritonymph (between the deutonymph and adult stages) are calyptostatic.

The more heavily-studied larval stages of the Trombiculoidea (commonly referred to as chiggers or berry bugs) primarily feed on terrestrial vertebrates, including humans [20], although some little-known taxa in this superfamily are ectoparasites of invertebrate hosts in common with the Trombidioidea [21-23] (Fig. 2). Importantly, the only major mite-transmitted disease of humans, scrub typhus or tsutsugamushi disease, is vectored by chiggers in the genus *Leptotrombidium* [24], while other chigger genera have only been implicated epidemiologically as locally-important vectors [25]. Scrub typhus is a severe febrile illness caused by infection with an obligate intracellular bacterium (*Orientia* spp.) in the order *Rickettsiales* and features an epidemiological cycle that includes wild small mammals, which are the primary hosts for many chigger species [26]. This disease has a fatality rate of 6% if not treated promptly with antibiotics [27] and has increased in incidence globally in recent years, with the annual minimum incidence reaching >17/100,000 in South Korea and Thailand in 2012 – 2013 [28]. In the so-called “tsutsugamushi triangle” within the Asia-Pacific region, scrub typhus has a median seroprevalence of 22.2% [28]; but endemic scrub typhus has emerged in several other parts of the world within the past decade, including South America [29], the Middle East [30], and possibly sub-Saharan Africa [31]. Chiggers have also been implicated in the transmission of hantaviruses [32], *Bartonella* spp. [33] and *Rickettsia* spp. [34]. Furthermore, chiggers have direct impacts worldwide by causing trombiculiasis, which is a highly pruritic dermatitis that can afflict humans, companion animals and domestic ruminants, potentially leading to severe hypersensitivity [35-38].

A remarkable second unique feature of trombidid mites is that the larvae induce the formation of a feeding tube or “stylostome” at the attachment site that is extraneous to the larval mouthparts [39]. These larvae are not blood feeders, but ingest tissue exudates (in the case of vertebrate hosts) or arthropod haemolymph [40]. The life history of trombidid nymphs and adults has been poorly studied. However, in the Trombiculoidea, the eggs of Collembola (springtails) and other arthropods are an important part of the diet [41] (Fig. 2). Arthropod eggs may also serve as food items for adults and nymphs in the Trombidioidea [42], although some species have potential roles in biological control, as they feed on pest arthropods such as spider mites, scale insects, aphids and termites [19, 43-45].

To date, research on trombidid mites has suffered from a dearth of molecular data that could facilitate studies on speciation; population structure; host-vector and vector-pathogen interactions; and life-history evolution in this group. To address this deficit, here we present a comparative analysis of the genomes of the chigger *Leptotrombidium deliense* (the primary scrub typhus vector in South-East Asia [46]) and the giant red velvet mite, *Dinothrombium tinctorium* (the world’s largest acarine species [47]). We show that these trombidid mites form a distinct branch of the Trombidiformes that exhibit two classes of LGT for secondary metabolism: the previously identified carotenoid biosynthesis enzymes of fungal origin, and much larger terpene synthase gene families, which probably derive from soil-associated bacteria. We also identify unique clusters of predicted allergens in *L. deliense* that may contribute to the symptoms of trombiculiasis in humans and domestic animals.

## Data description

Since the specimens were highly disparate in physical size (adult *D. tinctorium* can reach ~16 mm in length, whereas larval *L. deliense* do not normally exceed 250 μm, even when engorged), a tailored approach to sequencing was necessary in each case. For the velvet mite, DNA from a single adult was used to generate two Illumina TruSeq libraries (insert sizes, 350 bp and 550 bp) and one Nextera mate-pair library (insert size, 3 Kb). The TruSeq libraries were barcoded, indexed and paired-end (PE)-sequenced (2 × 100 bp) on one lane, and the Nextera library was PE-sequenced (2 × 250 bp) on an additional lane, both on the Illumina MiSeq platform. For the chiggers, DNA from a pool of engorged larvae (obtained from Berdmore’s ground squirrels in Thailand) was used to produce one NEB Next Ultra DNA library (insert size, 550 bp) and PE-sequenced (2 × 150 bp) on the Illumina MiSeq.

The total number of trimmed reads generated was ~362 million for *D. tinctorium* and ~38 million for *L. deliense*. For the former, PE reads were assembled using Abyss (v. 1.5.2) [48, 49]. For the chigger data, a preliminary assembly to contig level was performed using Velvet (v. 1.2.07) [50]. Reads derived from mammalian host genomic DNA were removed from the preliminary genome assembly using blobtools (v0.9.19), which generates a GC-coverage plot (proportion of GC bases and node coverage) [51] (Fig. 3). The *L. deliense* genome was then reassembled using SPAdes assembler (v. 3.7.1) [52] with default settings. For gene calling, the MAKER pipeline [53] was used to integrate *ab initio* gene predictions from Augustus (v. 3.2.2) [54], SNAP (v. 2013-11-29) [55] and GeneMark (v. 2.3e) [56] with evidence-based gene models. The “Methods” section provides more details on how downstream genome analyses were performed.

**Figure 3:**
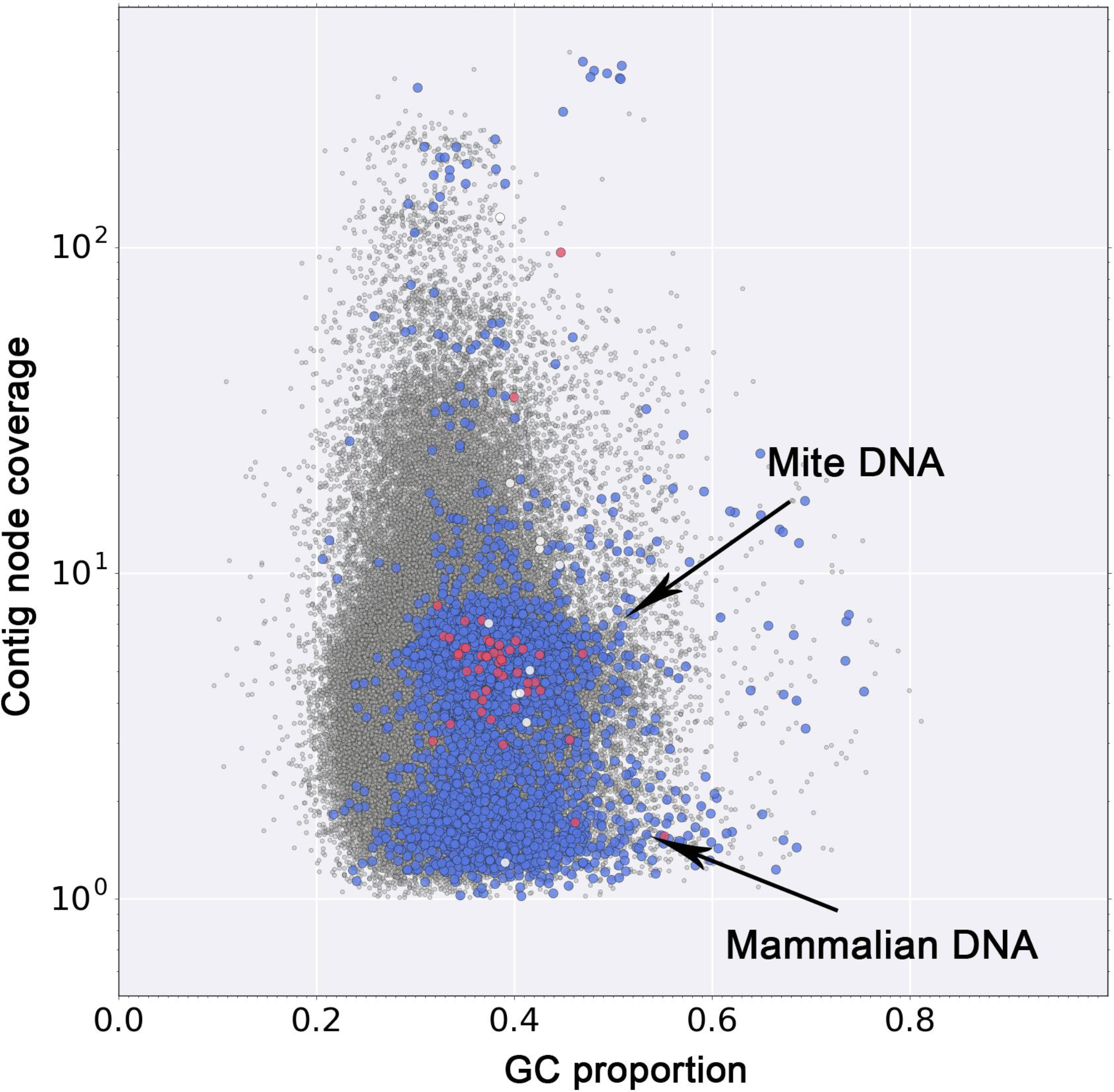
Blob-plot of contigs assembled from sequence data derived from engorged *Leptotrombidium deliense* larvae. Blue = Eukaryota; red = Bacteria; white = other; grey = no hit.

Similarly to the genome sequencing strategy, proteomic analysis of the trombidid mites was customised to the sample types available. A single adult *D. tinctorium* was subjected to protein extraction in SDS buffer and tryptic digestion using the filter-aided sample preparation method [57]. The digested sample was split into eight fractions using a High pH Reversed-Phase Peptide Fractionation Kit (Pierce) prior to nanoLC MS ESI MS/MS analysis on a Q-Exactive mass spectrometer (Thermo Fisher Scientific). A total of 137,638 spectra were generated across the eight fractions. For *L. deliense*, a soluble protein extract was obtained from a small pool of ethanol-fixed engorged larvae (n = 10) collected from several species of wild rodents in Thailand [26]. Following tryptic digestion, downstream analyses proceeded as for *D. tinctorium*, producing 18,059 spectra. The “Methods” section provides details on how MS spectra searches and Pfam enrichment analyses were performed.

## Analyses

### Genome statistics and phylogenomics

Assembled genome sizes were 180.41 Mb for *D. tinctorium* and 117.33 Mb for *L. deliense* (Additional file 1), whereas *k*-mer analysis placed the genome size estimates much closer together but was of a similar scale to the assemblies (143.52 – 147.09 and 158.31 – 160.95 Mb, respectively; Additional file 2). The repeat content of these genomes presented a significant challenge, with unclassified repeats alone accounting for 19 - 23% of the total size; approximately double the proportion of the *S. scabiei* [58] and *Tetranychus urticae* (two-spotted spider mite [59]) genomes (Additional file 3). As expected from sequencing a pool of *L. deliense* larvae compared with a single adult of *D. tinctorium*, and the corresponding DNA library strategies employed, the chigger genome was considerably less contiguous than that of the velvet mite. Nevertheless, the estimated completeness of the predicted gene set for *L. deliense* based on the Benchmarking Universal Single-Copy Orthologues (BUSCO) criteria [60] compared favourably with that of other arachnid genomes (Additional file 1). Notably, the *D. tinctorium* genome contained the greatest number of protein-coding genes from the Acariformes sequenced to date (Additional file 1).

Maximum likelihood and Bayesian phylogenomic analyses based on >500 one-to-one orthologous genes exhibited complete concordance, placing *L. deliense* and *D. tinctorium* together as sister taxa, and *T. urticae* as their closest relative amongst sequenced species (Fig. 1, Fig. 4). Our divergence time estimates accord closely with those previously published for arachnids [61], with the Parasitiformes and Acariformes separating approximately 416 million years ago (MYA) (Fig. 4). We estimate that the trombidid mites *sensu stricto* (velvet mites and chiggers) diverged from the phytophagous Tetranychoidea 250 MYA, and finally the Trombidioidea and Trombiculoidea last shared a common ancestor approximately 123 MYA (Fig. 4).

**Figure 4:**
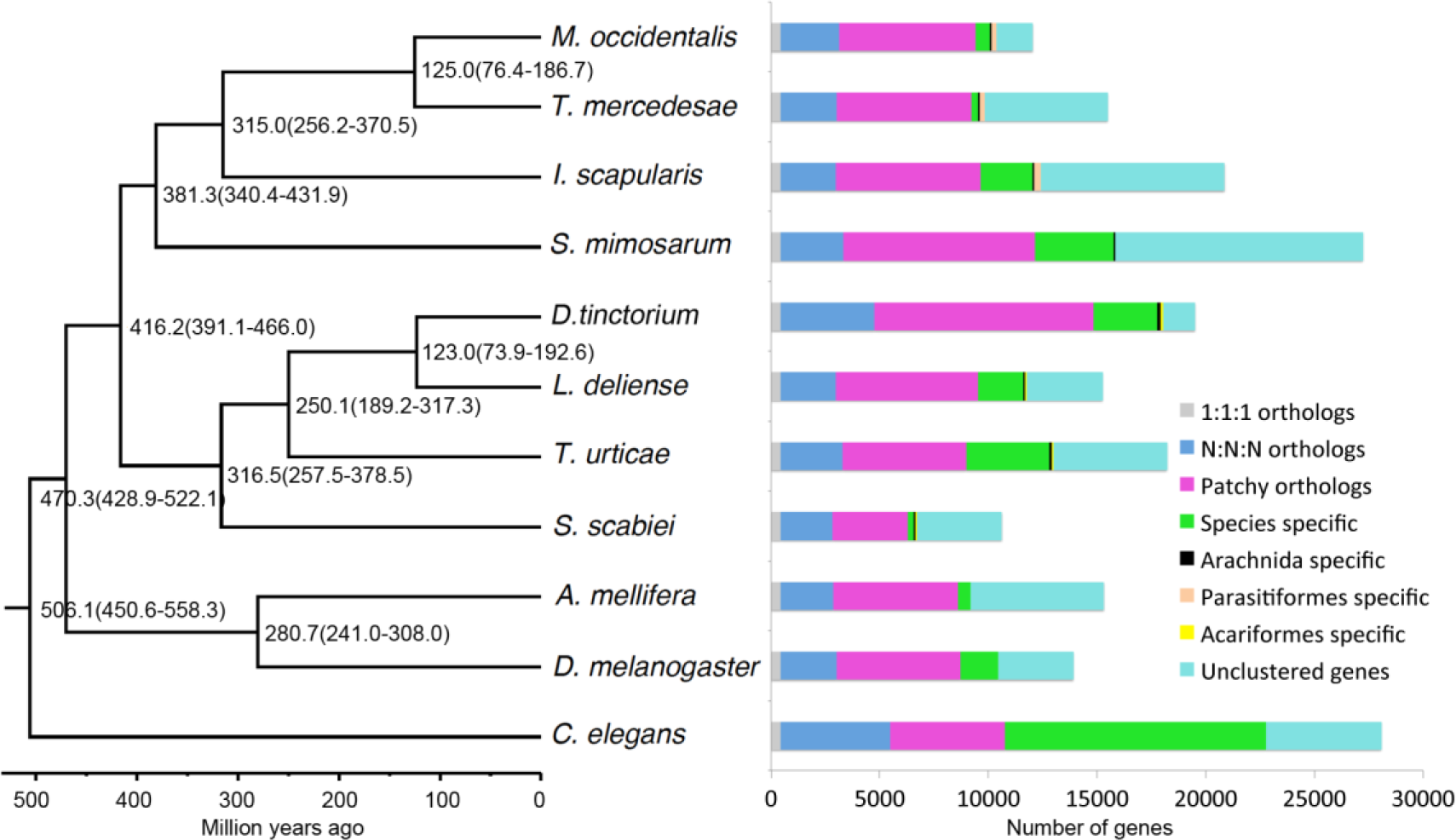
Estimated divergence times using a relaxed molecular clock with fossil calibration time and classification of protein-coding genes between 11 species of Ecdysozoa. *Caenorhabditis elegans* was used as the outgroup and the bootstrap value was set as 10,000,000. The 1:1:1 orthologs comprise the common orthologs with the same copy numbers in different species, and the N:N:N orthologs comprise the common orthologs with different copy numbers in these species. Patchy orthologs are shared between more than one, but not all species (excluding those belonging to the previous categories). Unclustered genes are those that cannot be clustered into gene families.

### Gene family expansions

When gene families were compared among the Acariformes and reference invertebrate genomes, the *D. tinctorium* genome was shown to contain a substantially greater number of unique paralogous groups than the other sequenced acariform mites, while only 68 gene clusters were shared among all Acariformes (Additional file 4). Moreover, the *D. tinctorium* genome displayed a greater number of multi-copy (“N:N:N”), patchy, and arachnid-specific orthologues when analysed alongside the published arachnid genomes (Fig. 4). The gene family expansion in *D. tinctorium* also dwarfed that seen among other members of the Arachnida, including *L. deliense* (Additional file 5).

Relative to other arachnids, *D. tinctorium* exhibited a significant expansion of 38 gene families, including a large family (ORTHOMCL98) of uncharacterised proteins containing 45 members in this species but no representatives in the other arachnid genomes (Additional file 6). Examination of conserved domains in this gene family revealed a major facilitator superfamily domain with some weak but significant similarity by BLASTP (~25% amino-acid identity, >95% coverage) to feline leukemia virus subgroup C receptor-related protein (FLVCR)-1 from various Metazoa. Two other orthologous clusters displayed an identity of up to 30% with FLVCR2 and showed a statistically significant expansion in the velvet mite, containing 27 (ORTHOMCL265) and 20 (ORTHOMCL412) members, compared with only one and three members in *L. deliense*, respectively (Additional file 6). Moreover, a gene family annotated as 4-coumarate:CoA ligases was significantly expanded in *D. tinctorium*, with 27 members; while *L. deliense* also showed a large repertoire (14 members) relative to other arachnids, although this was not statistically significant (Additional file 6). This gene family represents a group of long-chain fatty acyl-CoA synthetases related to firefly luciferase (but without luciferase activity) that are found in multicellular eukaryotes other than vertebrates [62]. These enzymes are associated with peroxisomes and perform catabolism of very long-chain fatty acids by β-oxidation.

Only a single gene, SPARC-related modular calcium-binding protein 1 (SMOC1) from *L. deliense*, was predicted to be under positive selection in either of the trombidid genomes with strong statistical confidence (dN/dS = 1.0663). The SMOC genes have been little studied in invertebrates, but in vertebrates they encode for matricellular proteins of the basement membranes and are involved in bone morphogenesis and ocular development [63]. The association with morphological development is conserved in *Drosophila*, in which the SMOC orthologue *pentagone* is expressed in the wing imaginal discs [64].

### Overrepresented protein families detected by LC-MS/MS

From the extremely small sample of engorged *L. deliense*, 522 mite proteins were identified by shotgun LC-MS/MS with at least one unique peptide against a background of 292 proteins of putative host origin (Additional file 7). Of the *L. deliense* proteins, 290 were considered as high-confidence identifications (≥2 unique peptides) and were subjected to Pfam domain enrichment analysis. The most overrepresented protein domains were derived from ATP synthase (PF00006 and PF02874) and both muscle and non-muscle isoforms of myosin or paramyosin (PF01576), although key enzymes of the citric acid cycle (ATP citrate synthase and succinate-CoA ligase, PF00549) were also well represented (Fig. 5). Several other proteins with probable origins from muscle tissue contained calponin homology (CH) domains (PF00307) and/or spectrin repeat domains (PF00435), including muscle protein 20 [65], myophilin, actinin, and spectrin subunits (Fig. 5). Myophilin is an invertebrate-specific protein that has previously been characterised as an immunogenic muscle component from parasitic platyhelminths [66], suggesting that it may contribute to the inflammatory response during trombiculiasis.

**Figure 5:**
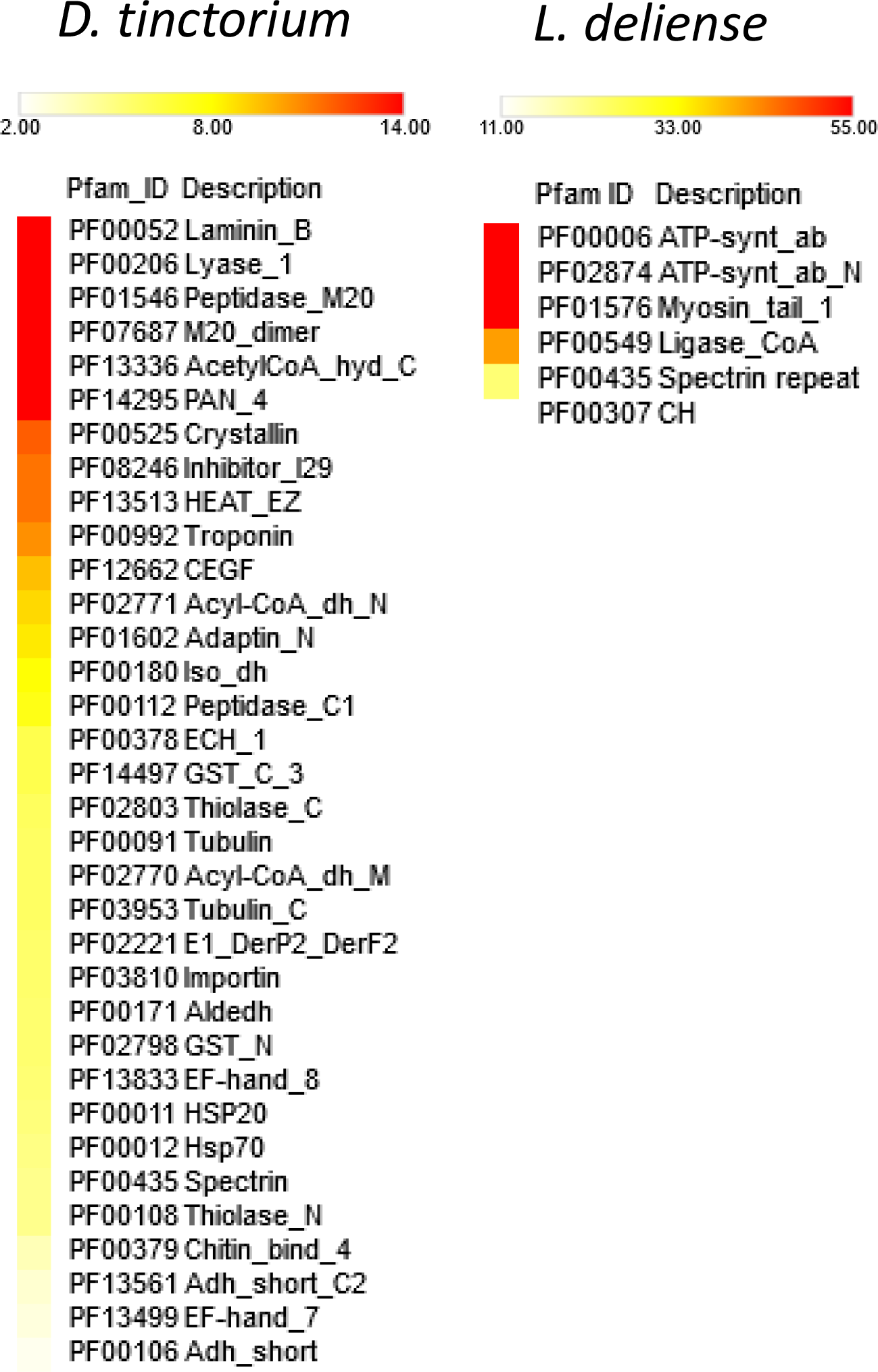
Overrepresented Pfam domains in proteomic datasets generated from a single adult *Dinothrombium tinctorium* and a pool of engorged *Leptotrombidium deliense* larvae. The colour scale represents fold-enrichment.

Proteomic analysis of a single adult *D. tinctorium* led to the robust identification (≥2 unique peptides) of 1,636 proteins, as the large size of the specimen was conducive to peptide fractionation prior to LC-MS/MS (Additional file 8). Laminin B (PF00052) was the most enriched domain (Fig. 5), which was present in several isoforms of perlecan (the basement membrane-specific heparan sulfate proteoglycan core protein). Perlecan is highly expressed during embryonic development in *Drosophila* [67], suggesting that this mite specimen may have contained fertilised eggs. Indeed, this specimen was undoubtedly a female, as it contained vitellogenins (yolk proteins) (Additional file 8). Lyase-1 domains (PF00206) were also highly enriched (Fig. 5) and were found in mitochondrial fumarate hydratase (an enzyme of the citric acid cycle) and in adenylosuccinate lyase of the purine nucleotide cycle. The peptidase M20 (PF01546) and M20 dimer domains (PF07687) were both present in cytosol non-specific dipeptidases, which have a key role in protein digestion in the midgut [68], and in N-fatty-acyl-amino acid synthase-hydrolases, which regulate thermogenesis via uncoupled respiration [69]. Another group of proteins with a putative role in thermoregulation were the α-crystallin-like small heat shock proteins (PF00525); in ticks, these are highly immunogenic proteins expressed in tick salivary glands and exhibit thermoprotective activity [70]. Finally, the inhibitor I29 domain (PF08246) represented digestive cysteine proteinases [71] and cathepsin L-like proteases (Fig. 5), which are highly expressed in feeding stages of *T. urticae* [72] and are a major protein component of spider mite faeces [73].

### Lateral gene transfers and mobile elements

To determine the origin of the bright colouration of the trombidid mites, we searched both genomes for the fused carotenoid synthases-cyclases that were reported to have been laterally transferred into the *T. urticae* genome from zycomycete fungi, perhaps via aphids [59]. In common with *T. urticae*, two of these carotenoid synthases-cyclases were observed in the *L. deliense* genome, while *D. tinctorium* harboured 12 copies (Additional file 9). However, the single largest gene family expansion observed in the *L. deliense* genome was within an orthologous cluster annotated as “pentalenene synthase”, which contained 41 members (Additional file 6). This cluster (ORTHOMCL47) also contained 21 genes in the *D. tinctorium* genome, but lacked orthologues in the genomes of other arachnids. A second orthologous cluster (ORTHOMCL881) of terpene synthases contained 17 members and was unique to *L. deliense* (Additional file 6).

The capacity to generate terpenoids (also known as isoprenoids) *de novo* in metazoans is extremely unusual. While some millipedes (for instance, the Japanese species *Niponia nodulosa*) are known to produce terpenes such as geosmin and 2-methylisoborneol in defensive secretions, these secondary metabolites are assumed to be derived from microbial symbionts [74]. The absence of terpene synthases in the fully sequenced diplopod genome from the rusty millipede, *Trigoniulus corallinus*, certainly supports this interpretation [75]. Dust mites also produce a monoterpene, neryl formate, which has been demonstrated to act as an aggregation pheromone [76]. However, BLAST analysis of the *Dermatophagoides farinae* genome assembly [77] using the trombidid terpene synthases failed to identify significant homologues, suggesting that dust mites rely on microbial symbionts for terpene production, or that terpene synthases in mite genomes are evolving too rapidly to be identified by homology searches. To the best of our knowledge, the only animals known to harbour terpene synthase genes in their nuclear genomes are a very restricted number of beetle species [predominantly flea beetles of the subfamily Galerucinae, which produce (6*R*,7*S*)-himachala-9,11-diene as a male aggregation pheromone [78]] and the collembolan *Folsomia candida*, in which the metabolites produced and their function are unknown [79]. Whilst terpene synthases are widespread in plants, fungi and bacteria, the terpene synthases from flea beetles do not resemble those from non-metazoan taxa and appear to have evolved from arthropod *trans*-isoprenyl diphosphate synthases [78]. Moreover, the terpene synthases of *F. candida* are more similar to those from the flea beetles than they are to the non-metazoan enzymes [79].

We generated phylogenies for the trombidid terpene synthases, which clearly showed close affinities to bacterial and fungal homologues and not to those from other arthropods (Fig. 6, Fig. 7). For ORTHOMCL47, the *L. deliense* and *D. tinctorium* enzymes formed distinct groups, and the closest homologues from other taxa included a monoterpene synthase from *Micromonospora* spp. (phylum Actinobacteria), a genus that is known to synthesise 2-methylenebornane [80], as well as related genes from agaricomycete fungi (Fig. 6). In the case of the chigger-specific ORTHOMCL881, the nearest homologues were distributed among Actinobacteria and other bacterial phyla, including the Chloroflexi, Proteobacteria and Bacteroidetes (Fig. 7). These terpene synthases were clearly separated from the flea beetle proteins and consisted predominantly of germacrene or geosmin synthases (Fig. 7). However, for both clusters of trombidid terpene synthases, the amino acid identity with their nearest bacterial or fungal homologues was low (≤30%). Nevertheless, the majority fulfilled the criteria of Crisp *et al.* [6] as high-confidence (“class A”) lateral gene transfers due to the absence of sufficiently closely-related homologues in other Metazoa.

**Figure 6:**
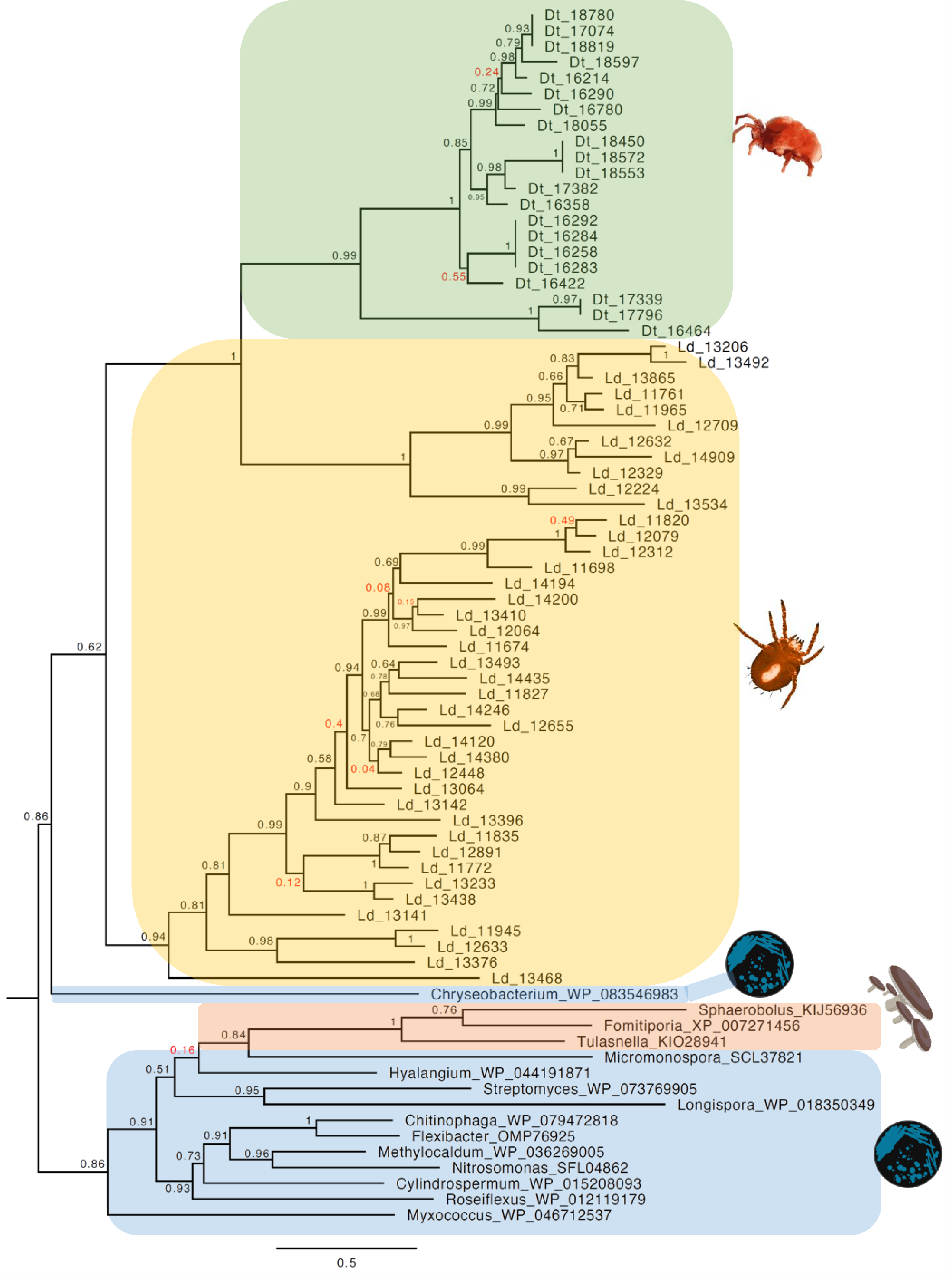
Phylogeny of terpene synthases from cluster ORTHOMCL47 of trombidid mites alongside related genes from bacteria and fungi. The tree was constructed using a maximum likelihood method; poorly-supported nodes are highlighted in red font.

**Figure 7:**
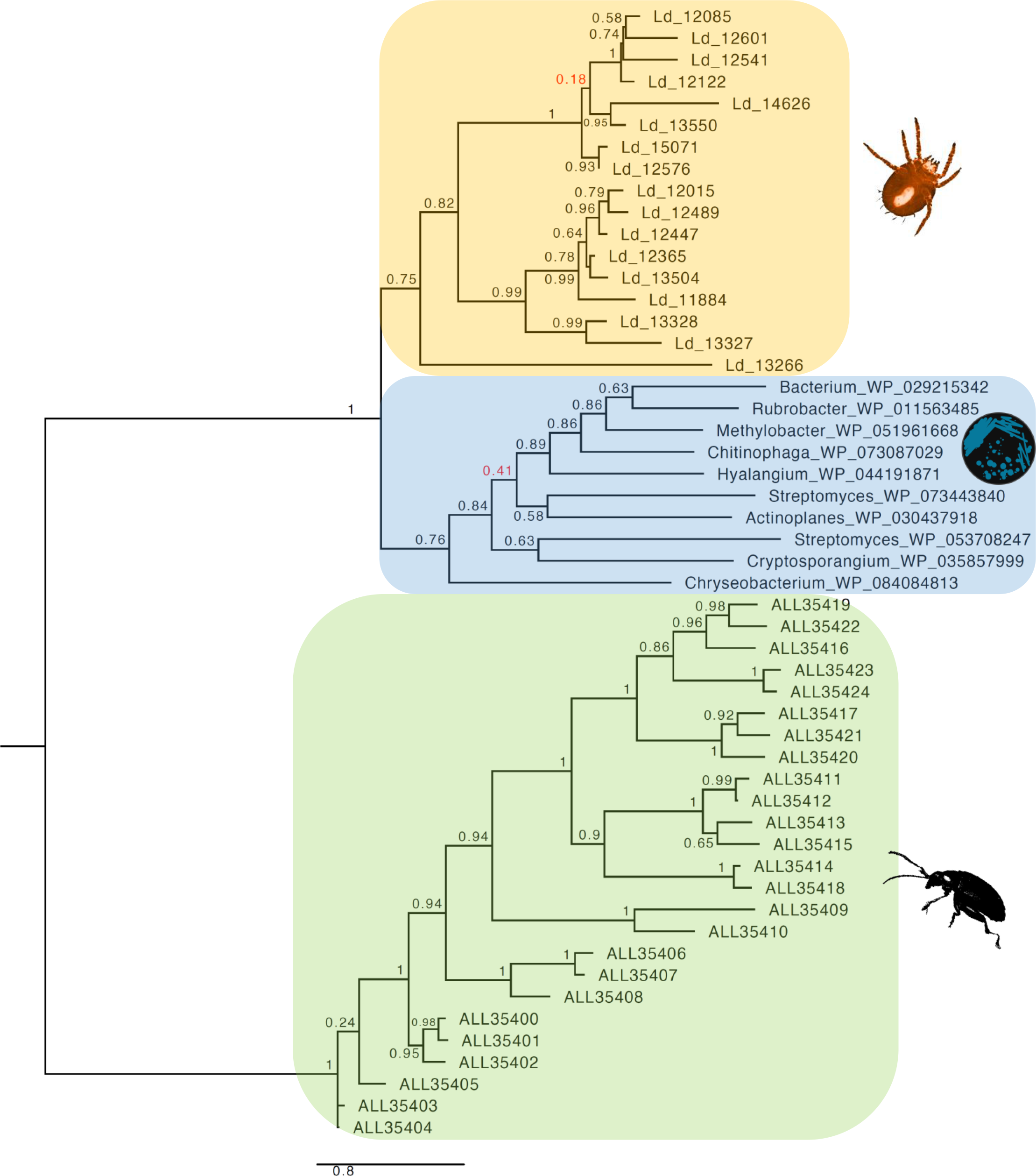
Phylogeny of terpene synthases from cluster ORTHOMCL881 of *Leptotrombidium deliense* alongside related genes from bacteria. Metazoan terpene synthases from galerucid beetles are shown as an outgroup. The tree was constructed using a maximum likelihood method; poorly-supported nodes are highlighted in red font.

To exclude the possibility that the trombidid terpene synthases were contaminating sequences of bacterial or fungal origin from the environment, or derived from microbial symbionts, we examined the genomic context of each terpene synthase to determine if they were sometimes found adjacent to an incontrovertible metazoan gene. Due to the low contiguity of the *L. deliense* assembly, it was not possible to find other genes on the same contig as a terpene synthase. However, one of the members of ORTHOMCL47 in *D. tinctorium* was located 5.2 kb downstream of a gene with a top BLAST hit to a translational elongation factor-2 mRNA from *Dinothrombium pandorae* [81] (Additional file 10). Furthermore, blob-plot analysis revealed that the GC content and read coverage of the contigs containing terpene synthases lay close to the overall mean for both trombidid genomes (Fig. 8). Importantly, sequences of unambiguous bacterial origin were very rare in both genomes (Additional file 11), with no evidence for a high-titre symbiont or environmental contaminant that may have impacted significantly on the genome assemblies. Of these candidate laterally-transferred genes in the trombidid mites, expression at the protein level could not be detected in the small chigger sample. However, in *D. tinctorium*, a high-confidence identification of a single terpene synthase from ORTHOMCL47 was achieved based on two unique peptides, while a third peptide was shared with an additional terpene synthase (Additional file 12).

**Figure 8:**
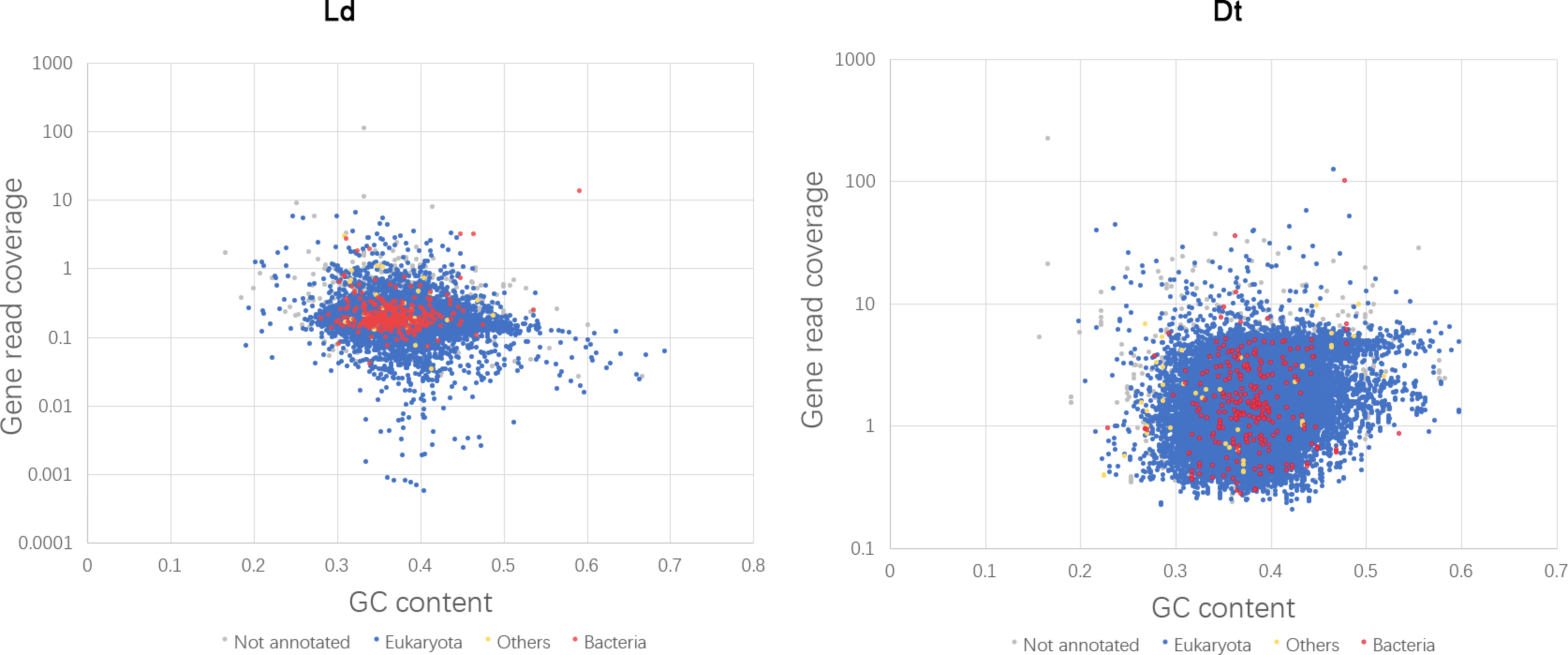
Blob-plot of read coverage and GC content for terpene synthase genes in *Dinothrombium tinctorium* (Dt) and *Leptotrombidium deliense* (Ld). Terpene synthases (in red, “Bacteria”) are shown in relation to all other mite genes (in blue).

In addition to these lateral gene transfers, the trombidid mite genomes exhibited further evidence for dynamism in the form of endogenous retroviruses (ERVs). The *D. tinctorium* genome showed significant expansions of reverse ribonuclease integrases and Pol polyprotein-like genes, whereas in the *L. deliense* genome, a 23-member family of Gag polyprotein-like genes was apparent (Additional file 6). Interestingly, the closest homologues of the Gag-like polyproteins in *L. deliense* were found in rodents, bats, lagomorphs, small carnivores, and colugos (a taxon restricted to South-East Asia [82]) (Fig. 9); all of which are known or likely hosts for chigger mites. Unfortunately, the low contiguity of the *L. deliense* genome and the metazoan context of ERVs (with similar GC content to the host) militated against bioinformatic attempts to exclude the possibility of an origin from host contamination; *i.e.*, from squirrel-derived cells on mite mouthparts or in the gut.

**Figure 9:**
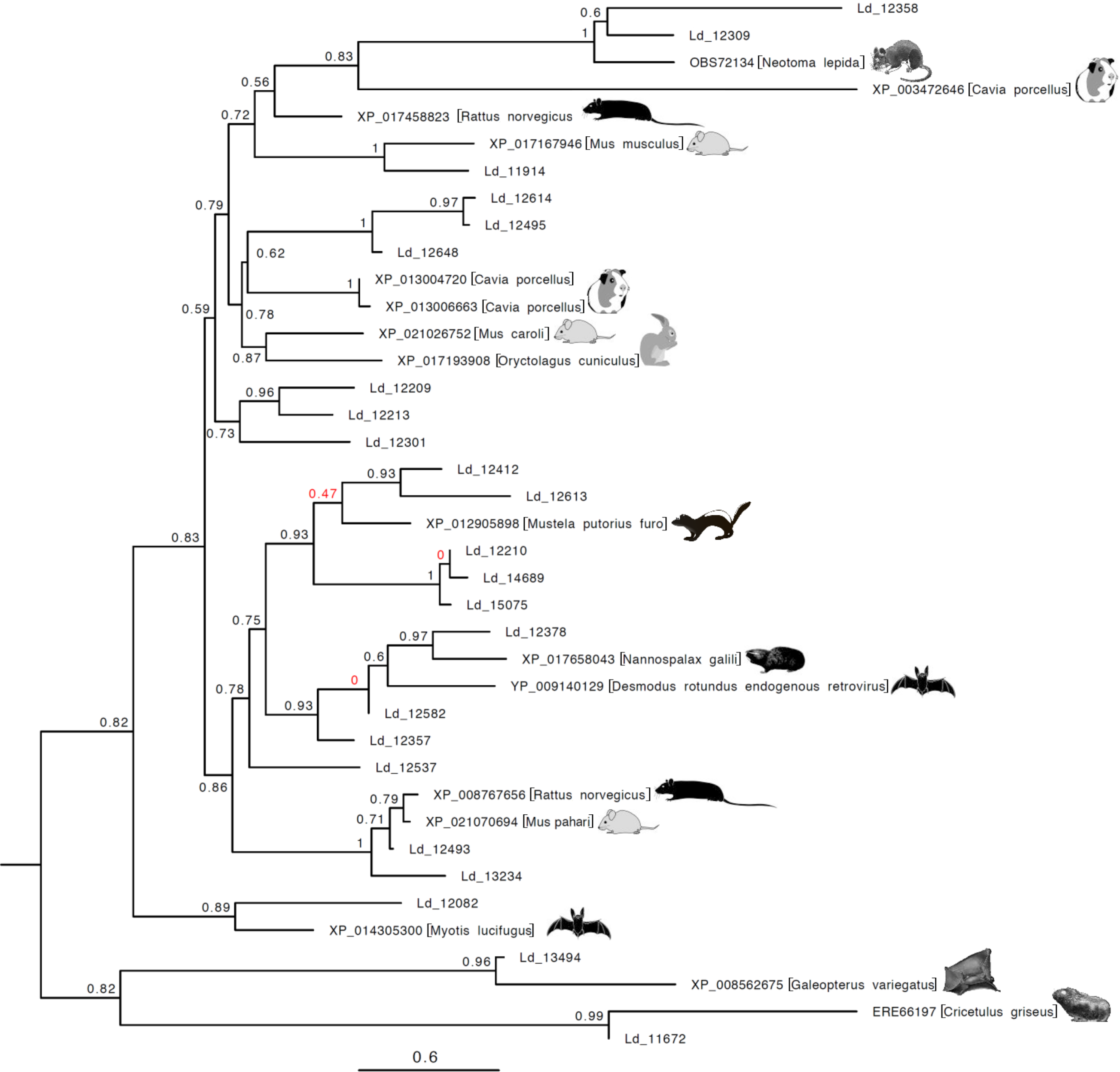
Phylogeny of Gag-like polyproteins from *Leptotrombidium deliense* in relation to homologous sequences from small mammals. The tree was constructed using a maximum likelihood method; poorly-supported nodes are highlighted in red font.

In the *D. tinctorium* genome, most members of the two ERV protein families each clustered in a monophyletic clade, suggesting expansion within the mite genome from a single origin (Additional files 13 and 14). The closest homologues of the Pol-like polyproteins were found mainly in other arthropods (especially cladocerans and ticks) and more distantly in fungi (Additional file 14); whereas the reverse ribonuclease integrases were most similar to those from ants, moths, nematodes and other mites (Additional file 13). Thus, endogenous retroviruses in *D. tinctorium* might originate from larval hosts, prey or soil microorganisms, although the fact that some velvet mites feed on other Acari [18] renders phylogenetic analyses of potential lateral gene transfers especially problematic. Uniquely among chelicerate genomes sequenced to date, the *D. tinctorium* genome also contained hepatitis D ribozyme-like genes (Rfam RF01787; Additional file 15), which in *Anopheles* mosquitoes, have been suggested to be involved in processing of non-LTR retrotransposons [83].

### Immune system

Many Acari act as vectors of plant or animal pathogens and their life histories expose them to a multitude of microorganisms in their diets and in the environment. Thus, how they interact with pathogens and commensals via their immune system is likely to be a critical aspect determining their success as a group. The canonical humoral immune response gene networks in *Drosophila* are the Toll signalling pathway (responding to β-1,3-glucans from fungi and lysine-type peptidoglycan from Gram-positive bacteria) and the immune deficiency (IMD) pathway (responding to diaminopimelic acid-type peptidoglycan from Gram-negative bacteria). These pathways are activated when upstream transmembrane receptors [peptidoglycan recognition proteins (PGRPs) and β-glucan recognition proteins] bind to the pathogen-derived molecules [84]. Recently, the expanding number of arthropod genomes from outside the class Insecta has highlighted key disparities in the immune pathway genes between the Pancrustacea (Hexapoda and Crustacea) *versus* the Chelicerata and Myriapoda. The most striking difference pertains to the IMD signalling pathway, which was thought to be absent in chelicerates [85]. However, genomic analyses and experimental data from *Ixodes scapularis* have revealed an alternative IMD pathway, in which interactions between IMD and Fas-associated protein with a death domain (both absent in the tick) are complemented by a E3 ubiquitin ligase (X-linked inhibitor of apoptosis protein, XIAP) and its ligand, the E2 conjugating enzyme Bendless [86]. This pathway recognises bacterial-derived lipids and restricts the growth of *Anaplasma phagocytophilum* and *Borrelia burgdorferi* in ticks. Although these data indicate that ticks (and perhaps other Parasitiformes) have a parallel IMD pathway distinct from that characterised in insects, we were unable to identify an XIAP homologue in acariform mites, including the trombidid genomes.

There are two non-exclusive scenarios that can be postulated to explain the apparent absence of an IMD pathway in acariform mites. The first is that these taxa might use the Toll pathway to respond to Gram-negative bacteria as well as Gram-positive bacteria and fungi. Indeed, crosstalk and synergistic immune responses to individual pathogens in *Drosophila* indicate that the two pathways are functionally interconnected even in insects [84], and the IMD pathway may have become redundant during the evolution of acariform mites. Second, expansions in other gene families associated with the immune response may provide alternative pathogen recognition and signalling pathways to tackle Gram-negative bacterial infections. This second scenario is supported in the trombidid mite genomes by large repertoires of Dscam genes (Additional file 16), which have previously been described to have undergone expansions in the Chelicerata and Myriapoda compared to the Pancrustacea [85]. In insects, Dscam is involved in phagocytosis of bacteria by haemocytes, and the *D. melanogaster* Dscam-hv gene exhibits a remarkable capacity to generate >150,000 alternatively-spliced isoforms, perhaps conferring some level of specificity to the insect immune response [although this remains highly controversial [87]]. Even relative to other acarine genomes, those of the trombidid mites display a substantially greater complement of Dscam genes (~40 in *D. tinctorium*; Additional file 16), rivalling the 60 gene family members observed in the *Strigamia maritima* (coastal centipede) genome [85, 88].

Several other expanded gene families in one or both of the trombidid mite genomes may have roles in the immune response. In common with *I. scapularis*, these genomes lack the transmembrane PGRPs that are activated in the presence of peptidoglycan in insects, but contain several other PGRP genes with putative extracellular or intracellular roles. However, these soluble PGRP genes are present in larger numbers in the trombidid mite genomes than in those of *I. scapularis* and *T. urticae* (Additional file 17). Since soluble PGRP fragments can have a co-receptor function as shown in insects [89], they might work in concert with as-yet-unidentified components of the acarine immune system to recognise pathogens. This is particularly important in the case of chiggers, as evidence for a peptidoglycan-like structure has recently been reported for *Orientia tsutsugamushi* [90]. Moreover, a much larger expansion in a second class of proteins with putative roles in the immune system, the C-type lectin domain (CTLD) proteins, was apparent in *L. deliense* (Additional file 6; Table 1). The CTLD protein family is a large and diverse group, most members of which do not bind carbohydrates and are thus not lectins [91]. If a CTLD protein does have lectin activity, the carbohydrate-recognition domain usually contains the amino acid motif “WND”, together with “EPN” if the specificity is for mannose, and QPD if the specificity is for galactose. However, several exceptions to this pattern do exist [91]. The expanded *L. deliense* CTLD proteins belong to four orthologous groups containing a total of 91 genes, of which one cluster (ORTHOMCL880) lacks any signatures of carbohydrate-binding activity (Table 1). The other three groups mainly contain proteins with EPN motifs, suggesting specificity for mannose, although a small proportion of QPD-motif CTLD proteins were apparent in two of the clusters, which might bind galactose (Table 1). The majority of the *L. deliense* CTLD proteins that were predicted to bind carbohydrates exhibited classical or internal secretion signatures, while only a small proportion (10 – 20%) contained transmembrane domains (Table 1). In common with many members of the CTLD protein family, including those in other arthropods, *N*-glycosylation sites were predicted in 10 – 20% of the *L. deliense* CTLD proteins [92] (Table 1).

**Table 1:**
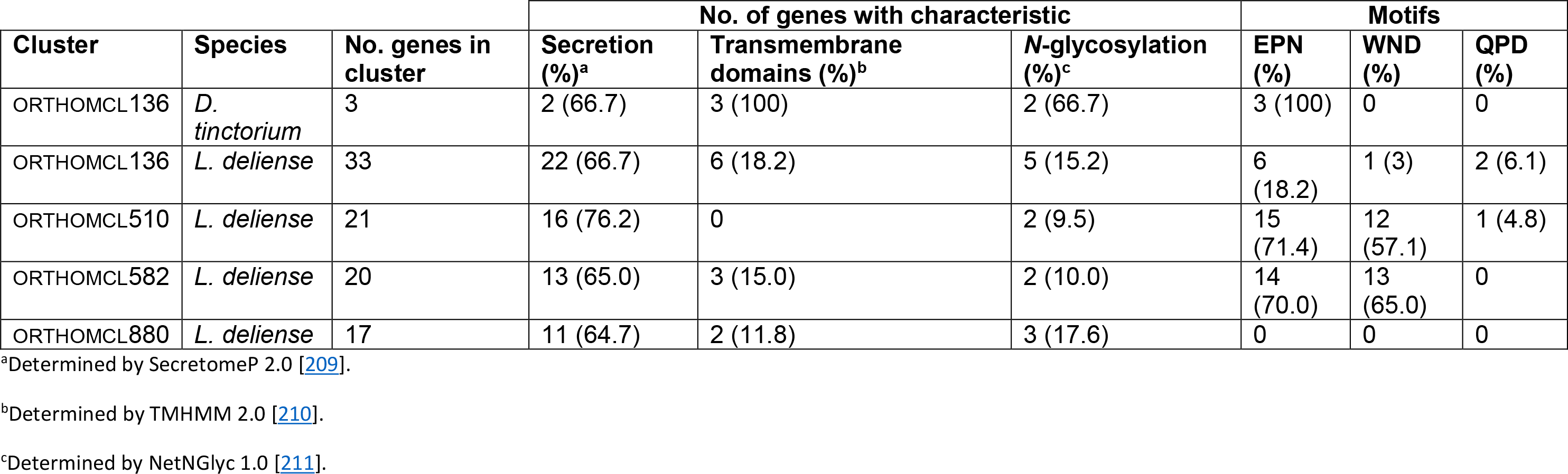
Characteristics of C-type lectin domain proteins in the trombidid mite genomes.

A final orthologous cluster with a potential role in innate immunity and which has undergone a significant expansion in *L. deliense* was annotated as “double-stranded (ds)RNA-activated kinase” (Additional file 6) or protein kinase R (PKR), an interferon-inducible enzyme that can block viral protein synthesis following binding to dsRNA substrates [93]. However, canonical PKR is restricted to mammals [94], and the 13 proteins from this cluster in the *L. deliense* genome lack a dsRNA-binding domain (InterPro identifier: IPR014720). Unexpectedly, the closest homologues to the PKR-like proteins in *L. deliense* were not from other arthropods but from a variety of different eukaryotic taxa, including protists and fungi (Additional file 18). However, the sequence similarity between the *L. deliense* genes and homologues in other Metazoa (principally in cnidarians and mammals) was sufficient to prevent fulfilment of the Crisp *et al.* [6] criteria for lateral gene transfer. Notwithstanding the absence of RNA-binding domains, these *L. deliense* proteins resemble PKR-like endoplasmic reticulum kinase (PERK), widespread in eukaryotes, which is a key component of the unfolded protein response during periods of endoplasmic reticulum stress [95].

### Photoreceptor and chemosensory systems

Unlike insects, chelicerates lack compound eyes. Mites and ticks may be eyeless, or can possess one or more pairs of simple dorsal ocelli. The Parasitiformes sequenced to date are all eyeless species (*I. scapularis, M. occidentalis* and *T. mercedesae*), whereas the trombidid mites and *T. urticae* have two pairs of ocelli on the prodorsum in the adult stage ([47]. However, the genomes of both eyeless and eyed Acari exhibit a variable complement of opsins, which in combination with the chromophore retinal, form light-sensitive proteins termed rhodopsins. The genomes of eyeless ticks and mites, as well as that of *T. urticae*, contain one or more genes of the “all-trans-retinal” peropsin class, which in spiders have been shown to encode non-visual photosensitive pigments with combined G-protein coupled receptor and retinal photoisomerase activity [96]. Since even the eyeless species show evidence for reproductive and diapause behaviours that respond to day-length, it has been suggested that peropsins are important for the maintenance of circadian rhythms [97, 98]. Notably, we found no evidence of peropsin genes in the trombidid mite genomes, but did find orthologues of *T. urticae* rhodopsin-1 and −7 in both *L. deliense* and *D. tinctorium* (Fig. 10). In the velvet mite, an additional four rhodopsin-7-like paralogues were apparent, three of which were identical at the amino-acid level (Fig. 10).

**Figure 10:**
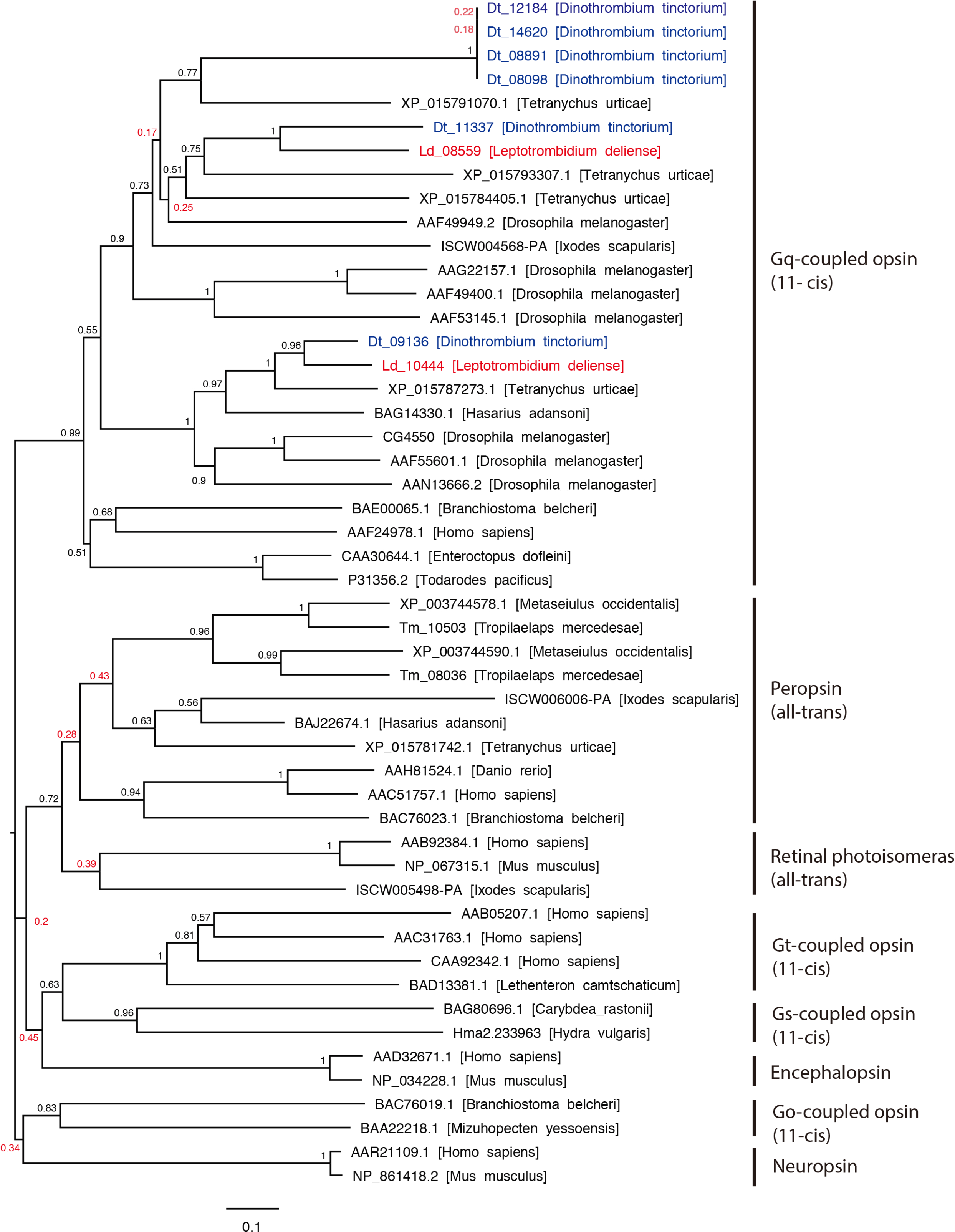
Phylogeny and classification of metazoan opsins. The tree was constructed using a neighbour-joining method. Poorly-supported nodes are highlighted in red font.

In contrast with insects but in common with crustaceans and myriapods, the Acari appear to have a scant repertoire of chemosensory protein classes, lacking both odorant-binding proteins (OBPs) and odorant receptors. Moreover, the small chemosensory proteins that have expanded considerably in some insect orders (especially Lepidoptera [99]) are completely absent in the mite genomes, although a gene encoding one such protein was identified in the *I. scapularis* genome (Table 2). Thus, mites rely primarily on gustatory and ionotropic receptors for chemosensation. The repertoire of gustatory receptors (GRs) in *L. deliense* (42 members) and *D. tinctorium* (105 members) was in a similar range to most mites and ticks (albeit from the Parasitiformes) and for the Mandibulata (Table 2); hence, there was no evidence for the massive expansion in this gene family recently reported for the *T. urticae* genome, with almost 700 members [100].

**Table 2:** Comparison of chemosensory receptor repertoires between trombidid mites and 11 other arthropods.

| Species | Chemosensory receptor<sup>a</sup> |  |  |  |  |
| --- | --- | --- | --- | --- | --- |
|  | GR | OR | IR | OBP | CSP |
| *D. tinctorium* | 105 | 0 | 7 | 0 | 0 |
| *L. deliense* | 42 | 0 | 8 | 0 | 0 |
| *T. urticae* | 689 | 0 | 4 | 0 | 0 |
| *T. mercedesae* | 5 | 0 | 8 | 0 | 0 |
| *M. occidentalis* | 64 | 0 | 65 | 0 | 0 |
| *I. scapularis* | 60 | 0 | 22 | 0 | 1 |
| *S. maritima* | 77 | 0 | 60 | 0 | 2 |
| *D. pulex* | 53 | 0 | 85 | 0 | 3 |
| *D. melanogaster* | 73 | 62 | 66 | 51 | 4 |
| *A. mellifera* | 10 | 163 | 10 | 21 | 6 |
| *B. mori* | 56 | 48 | 18 | 44 | 18 |
| *A. pisum* | 53 | 48 | 11 | 15 | 13 |
| *P. humanus*<br>*humanus* | 8 | 10 | 12 | 5 | 7 |

<sup>a</sup>GR, gustatory receptor; OR, olfactory receptor; IR, ionotropic receptor; OBP, odorant-binding protein; CSP, chemosensory protein. Data for other genomes were obtained from [88, 97, 98, 100, 213].

Ionotropic glutamate receptors (iGluRs) are glutamate-gated ion channels that are divided into two subtypes based on sensitivity to N-methyl-D-aspartic acid (NMDA). The canonical iGLuRs do not have direct roles in chemosensation. Rather, at least in *D. melanogaster*, the NMDA-sensitive channels are expressed in the brain and are involved in associative learning and memory [101]. The non-NMDA channels have fundamental roles in synaptic transmission in the neuromuscular junction within muscle tissue or in the nervous system [102], and certain receptor subunits have been shown to be involved in the regulation of sleep (GluR1 [103]) or vision (Clumsy, CG5621, CG3822, CG11155, CG9935 [104, 105]). Strikingly, the *D. tinctorium* genome harboured seven NMDA-type iGluRs and 61 non-NMDA iGluRs, representing substantially greater repertoires than those observed for the *L. deliense, T. urticae* and *D. melanogaster* genomes (especially for the non-NMDA iGluRs) (Fig. 11). The chemosensory ionotropic receptors (IRs), which exhibit sequence similarity to iGluRs but do not bind glutamate [106], also showed interesting differences in gene family size compared with *T. urticae* and *D. melanogaster*. Notably, while *D. melanogaster* has one gene encoding an IR25a protein, *T. urticae* has three such genes and the trombidid mites have five copies each (Fig. 11). The *D. melanogaster* IR25a is a widely-expressed co-receptor that couples with stimulus-specific IRs to facilitate sensitivity to a diverse range of acids and amines. Recently, IR25a in combination with IR21a and IR93a were demonstrated to function as a thermosensory complex expressed by the dorsal organ cool cells of *D. melanogaster* larvae, which mediates avoidance behaviour to cool temperatures (<20°C) [107, 108]. Sequences that cluster with *D. melanogaster* IR21a and IR93a in the “antennal and first leg” class of IRs were identified in the trombidid mite genomes, with one copy in *D. tinctorium* (as for *T. urticae*) and three copies in *L. deliense* (Fig. 11). Although chelicerates lack antennae, the orthologues of IR93a and/or IR25a have been shown to be highly expressed exclusively in the first pair of legs in *T. mercedesae* [98] and *Varroa destructor* [109], suggesting functional parallels between insects and mites.

**Figure 11:**
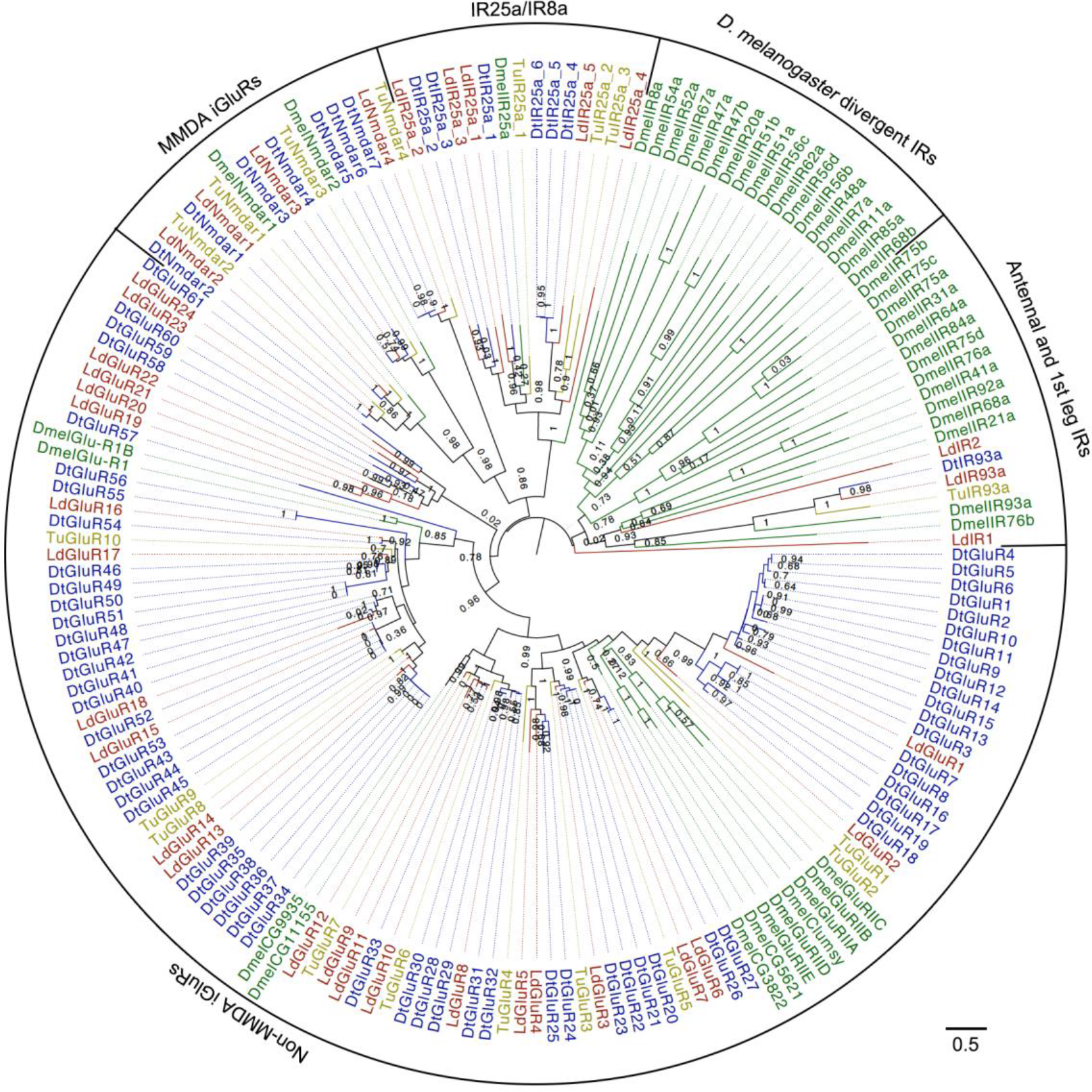
Phylogeny of *Dinothrombium tinctorium, Leptotrombidium deliense, Tetranychus urticae* and *Drosophila melanogaster* ionotropic receptors and ionotropic glutamate receptors. The tree was constructed using a maximum-likelihood method.

### Predicted allergens

Although the propensity of chiggers to cause pruritic dermatitis is well recognised in humans and other animals [35-38], the identity of the allergens involved has not been established [110]. The *L. deliense* and *D. tinctorium* genomes were predicted to encode 37 and 33 groups of protein allergens, respectively; substantially more than other sequenced mites in the Acariformes with the exception of the dust mites, *D. farinae* [77] and *E. maynei* [111]. Since velvet mites rarely come into contact with humans, only the chigger allergens were subjected to further analysis. The *L. deliense* predicted allergen clusters included nine groups that were unique to this species and six that were shared with *D. tinctorium* only (Fig. 12), while a further 28 putative allergen genes in the *L. deliense* genome did not cluster in orthologous groups (Additional file 19). These chigger-unique groups included five distinct clusters of trypsin-like serine proteases and one cluster each of subtilases, papain-like cysteine proteases, enolases, and cyclophilins, all of which could be classified into recognised allergen families listed in the AllFam database [112] (Fig. 13). The non-clustered allergens belonged to a variety of structural and enzymatic protein groups, but cathepsins, serine proteases and peptidylprolyl isomerases were the most common annotations (Additional file 19).

**Figure 12:**
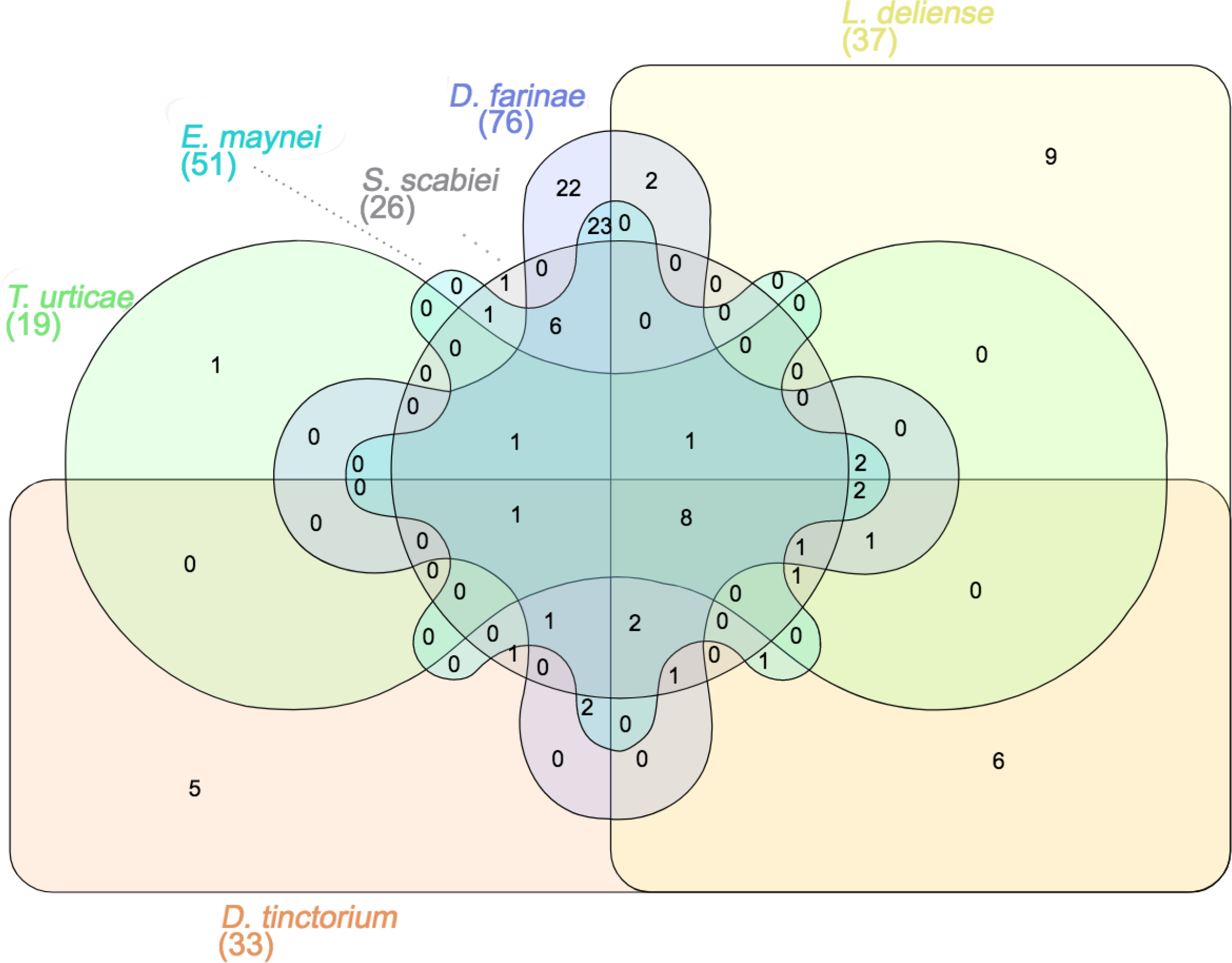
Venn diagram of orthologous clusters of predicted allergens from six species of acariform mites.

**Figure 13:**
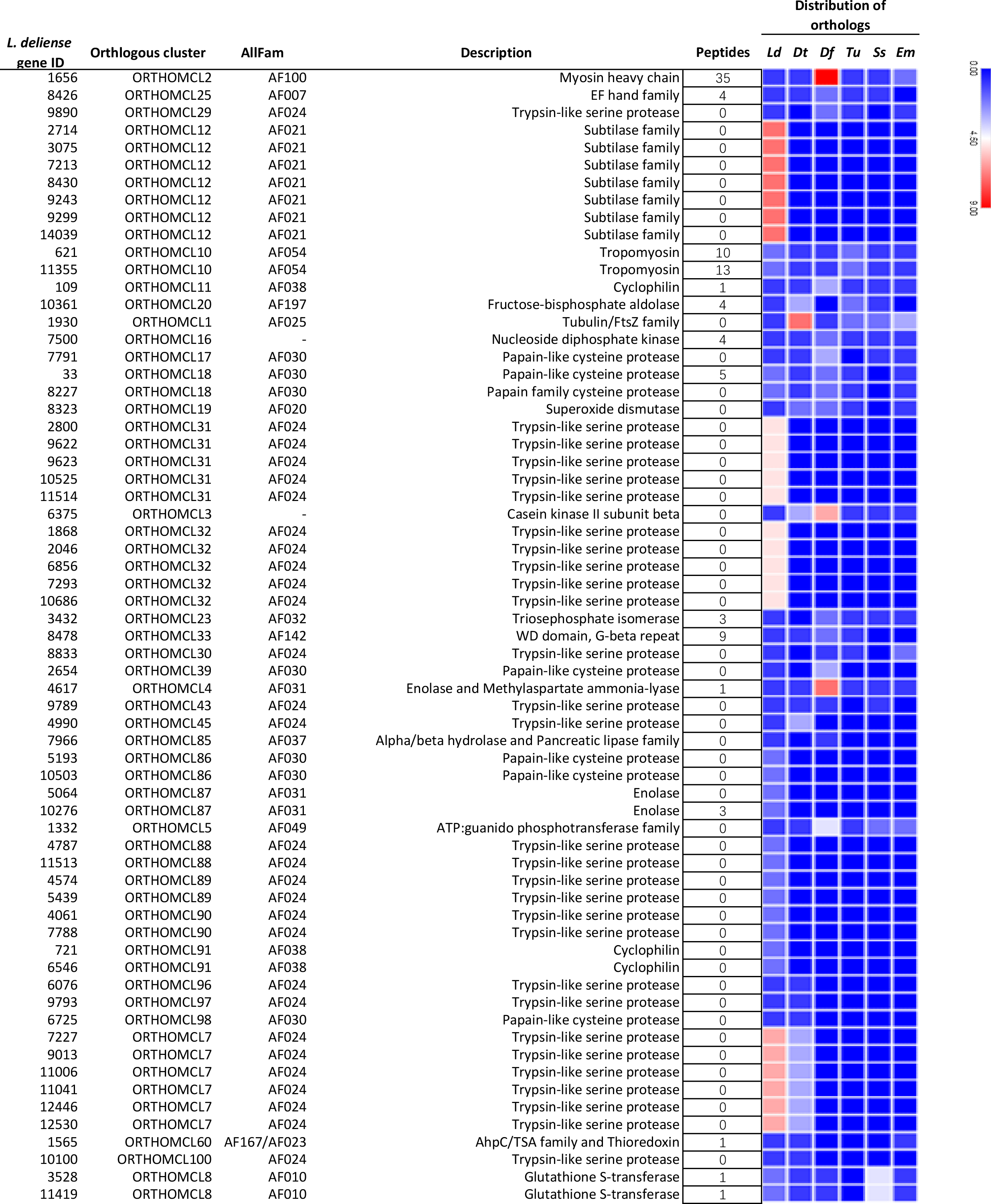
Orthologous clusters of predicted allergens in *Leptotrombidium deliense* as classified by the AllFam database. “Peptides” refers to the number of unique peptides from each allergen detected by mass spectrometric analysis of a pool of *L. deliense* larvae. The heat-map indicates the number of orthologues for each predicted allergen in the genomes of six acariform mites. Ld, *Leptotrombidium deliense*; Dt, *Dinothrombium tinctorium*; Df, *Dermatophagoides farinae*; Tu, *Tetranychus urticae*; Ss, *Sarcoptes scabiei*; Em, *Euroglyphus maynei*.

The major allergens in *D. farinae* are the 25-kDa Der f 1, a papain-like cysteine protease; and Der f 2, a 14-kDa uncharacterised protein with a ML (lipid-binding) domain [77]. However, many other minor allergens have been detected by immunoproteomic studies [113, 114] or predicted by homology searches in the *D. farinae* genome [77]. In *L. deliense*, five distinct clusters of papain-like cysteine proteases were identified (AllFam AF030), of which three were shared with *D. farinae*, one was shared only with *D. tinctorium*, and one was unique (Fig. 13). No orthologue of Der f 2 (AF111) was apparent.

Recently, an alpha-enolase has been reported as a novel minor allergen in *D. farinae* [114]. However, the two enolases (AF031) with predicted allergenic properties in the *L. deliense* genome formed an orthologous cluster that was absent from other mites sequenced to date, with homologues in parasitic nematodes and distant chelicerate relatives (*e.g.*, horseshoe crabs; Additional file 19). A similar pattern was observed for the cyclophilins (AF038), which have previously been considered a class of allergens restricted to fungal and plant sources [115], although these peptidyl-prolyl *cis-trans* isomerases are universally present across all domains of life. In an immunoproteomic study, a cyclophilin was newly identified as a dust mite allergen, Der f 29 [113], but this was not closely related to the *L. deliense* cyclophilins, which exhibited a greater affinity (~75% identity) to homologues in fungi and fish (Additional file 19). The *L. deliense* subtilases (serine proteases with a peptidase S8/S53 domain, AF021) were also absent from other mite genomes but showed 40 – 50% identity to subtilases from fungi and bacteria (Additional file 19). This class of proteases have been identified as major allergens produced by ascomycete fungi such as *Curvularia lunata* [116] and *Trichophyton* spp. [117]. Finally, the five clusters of trypsin-like serine proteases (AF024) exhibited closest homologues (40 – 50% identity) in a diverse range of organisms, including *T. urticae* (but too distant to cluster in ORTHOMCL31), Diptera and scorpions (ORTHOMCL32), fish and lizards (ORTHOMCL88), bugs and ants (ORTHOMCL89), and acorn worms and Diptera (ORTHOMCL90) (Additional file 19). Thus, these predicted allergens were distinct from the *D. farinae* molecules classified in AF024 [Der f 3, 6 and 9 [118]], although within ORTHOMCL29, ORTHOMCL30, and ORTHOMCL43, *L. deliense* does possess additional trypsin-like proteases that are orthologous to these *D. farinae* allergens (Figure 13).

A label-free quantitative analysis of protein content in the chiggers indicated that muscle-derived allergens related to the *D. farinae* paramyosin Der f 11 (AF100 [119]) and to *T. urticae* tropomyosin isoforms (AF054) were most abundant (Fig. 13). Although single unique peptides were detected for several *L. deliense-specific* allergen clusters and unclustered allergenic proteins, only one *L. deliense-*specific allergen was present in quantifiable amounts (an enolase in AF031), and this was considerably less abundant than the shared allergens (Additional file 7, Fig. 13). However, as allergenicity is not dictated entirely by allergen quantity and can vary markedly between individuals, validating the identity of the most important allergens in chiggers will require screening of sera from trombiculiasis patients.

### Putative salivary proteins

Due to the diminutive size of chiggers and the absence of any artificial feeding mechanism for laboratory colonies that might allow collection of saliva, the chigger sialome has not been characterised to date. However, numerous high-throughput studies of tick saliva have been conducted on several genera and multiple lifecycle stages [120-125], and recently an elegant proteomic analysis of *T. urticae* saliva was published, in which mite salivary secretions were collected in an artificial diet substrate [126]. Using proteomic datasets from this *T. urticae* study and a recent *I. scapularis* sialome analysis conducted over several time-points [120], we identified one-to-one orthologues of the salivary proteins from both sources in the tick, spider mite, and trombidid mite genomes. We reasoned that as *T. urticae* is phylogenetically close to the trombidid mites while *I. scapularis* is very distant, protein families shared by the tick and the trombidid mites but not present in the *T. urticae* genome are likely to represent proteins required for ectoparasitism on animal hosts (as opposed to phytophagy). Indeed, 24 orthologous clusters were shared among the animal ectoparasites but not with *T. urticae*, whereas only five clusters were shared between all mites at the exclusion of the tick (Figure 14). These 24 animal-ectoparasite clusters are candidates as key salivary components of trombidid mites. An additional two clusters were shared exclusively by *I. scapularis* and *L. deliense*, suggesting that they might be important for feeding on vertebrate hosts (Fig. 14).

**Figure 14:**
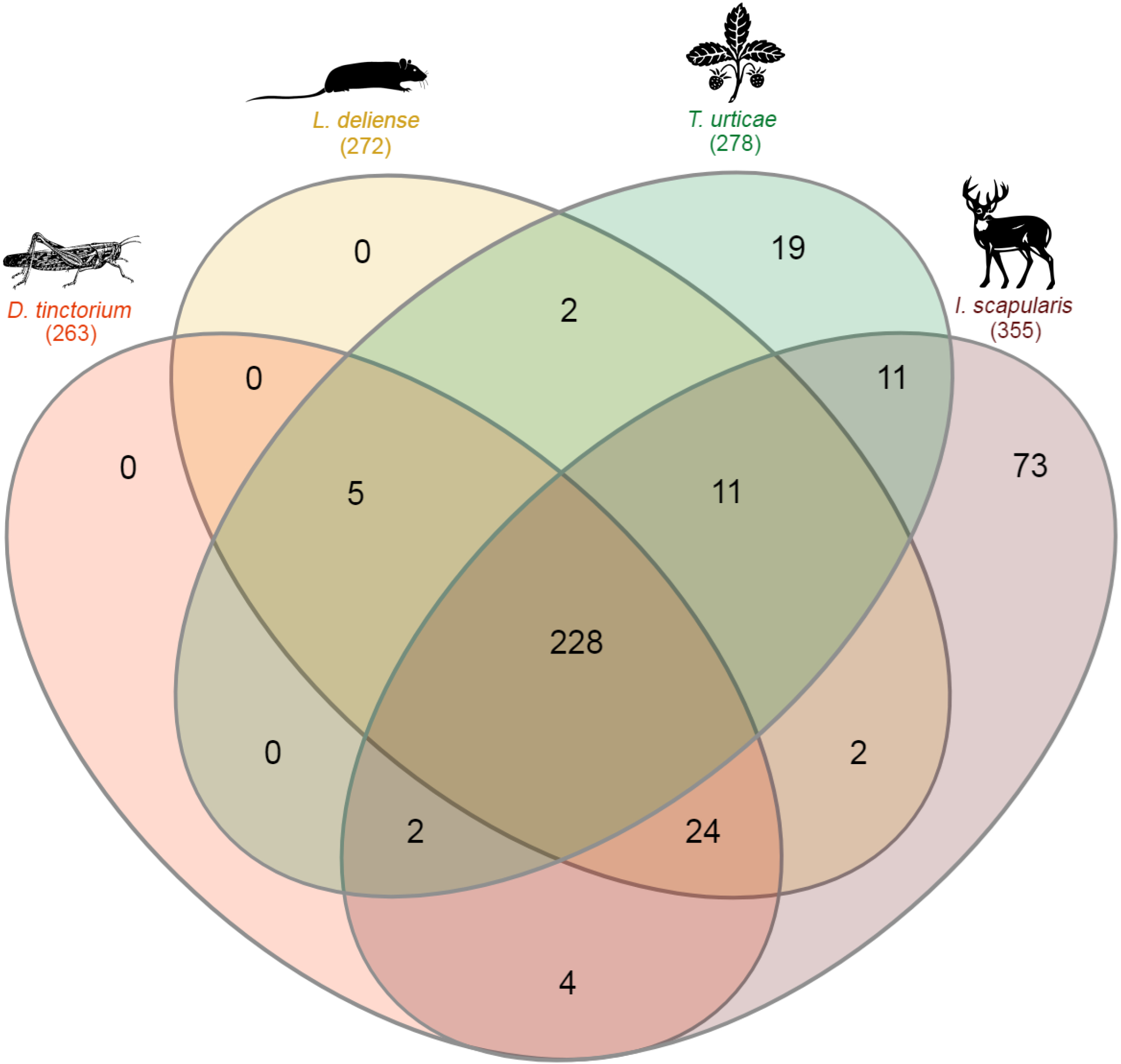
Venn diagram of orthologous clusters of putative salivary proteins in four species of Acari. One-to-one orthologues of salivary proteins from *Ixodes scapularis* (Kim et al.) and *Tetranychus urticae* (Jonckheere) were identified in the genomes of *Leptotrombidium deliense* and *Dinothrombium tinctorium*. Thumbnail images indicate representative host species.

To feed successfully, ticks must suppress local immune responses and prevent the clotting of blood. Although trombidid mites feed on tissue exudates or haemolymph rather than blood, and do not feed for as long as some hard tick species, they face similar challenges as ectoparasites that provoke an inflammatory response in their hosts. Interestingly, in accordance with low levels of haem in the trombidid mite diet, we did not find orthologues of tick salivary proteins involved in haem detoxification (ferritins and hemelipoproteins [122]) in the trombidid mite genomes. Several lipocalins with histamine-binding activity have been identified in tick saliva from multiple different species [120, 122, 123], but orthologues of these small proteins were also not present. However, genes encoding two enzymes involved in catabolism of the histamine precursor histidine, urocanate hydratase and formiminotransferase-cyclodeaminase, were detected in both trombidid mite genomes (Table 3). The degradation of histidine feeds into the one carbon pool by folate, and this process is mediated in part by formyltetrahydrofolate dehydrogenase, an enzyme that is also present in multiple copies in the *D. tinctorium* genome (Table 3). While the presence of folate biosynthesis enzymes in tick saliva has been reported previously (and not only for *I. scapularis* [122]), the functional significance of their secretion is unclear. One possibility is that ectoparasitic Acari not only utilise bacterial symbionts as folate “factories” [127, 128], but can scavenge it at source from precursors in their B-vitamin-deficient diets.

**Table 3:** Orthologues of *Ixodes scapularis* salivary proteins in trombidid mite genomes.

| Orthologous cluster | Number of genes in cluster |  |  | Representative gene ID | Representative annotation<sup>a</sup> |
| --- | --- | --- | --- | --- | --- |
|  | *I. scapularis* | *L. deliense* | *D. tinctorium* |  |  |
| ORTHOMCL5799 | 1 | 2 | 1 | EEC06447 | CNDP dipeptidase |
| ORTHOMCL8488 | 1 | 1 | 1 | EEC03480 | Uncharacterized protein (peptidase M17, leucine aminopeptidase/peptidase B domain)] |
| ORTHOMCL9 | 73 | 2 | 2 | EEC08659 | Sulfotransferase |
| ORTHOMCL2940 | 2 | 1 | 2 | ISCW018873 | Short-chain alcohol dehydrogenase |
| ORTHOMCL8622 | 1 | 1 | 1 | JAA64643 | Glycine-rich cell wall structural protein, partial [*Rhipicephalus pulchellus*] |
| ORTHOMCL3018 | 1 | 1 | 3 | EEC14134 | Calponin |
| ORTHOMCL5249 | 2 | 1 | 1 | JAB84323 | Stretch regulated skeletal muscle protein [*Ixodes ricinus*] |
| ORTHOMCL7670 | 1 | 1 | 1 | EEC14106 | Multifunctional chaperone |
| ORTHOMCL8293 | 1 | 1 | 1 | EEC09272 | Uncharacterised protein (aspartate dehydrogenase domain) |
| ORTHOMCL262 | 5 | 5 | 5 | ISCW002881 | UDP-sugar hydrolase |
| ORTHOMCL2418 | 1 | 1 | 4 | EEC13206 | Cytochrome C |
| ORTHOMCL5815 | 2 | 1 | 1 | ISCW011750 | Succinyl-CoA ligase beta subunit |
| ORTHOMCL3911 | 1 | 1 | 3 | JAB73948 | Formyltetrahydrofolate dehydrogenase, partial [*Ixodes ricinus*] |
| ORTHOMCL3015 | 2 | 1 | 2 | ISCW001951 | ATPase inhibitor |
| ORTHOMCL8 | 66 | 12 | 7 | EEC10817 | Acyl-CoA synthetase |
| ORTHOMCL260 | 13 | 1 | 1 | ISCW001079 | Acetylcholinesterase |
| ORTHOMCL6164 | 2 | 1 | 1 | ISCW022662 | NS1-binding protein |
| ORTHOMCL4904 | 2 | 1 | 1 | ISCW006538 | 60S acidic ribosomal protein LP1 |
| ORTHOMCL907 | 6 | 1 | 1 | EEC05404 | Translation initiation inhibitor UK114/IBM1 |
| ORTHOMCL4900 | 2 | 1 | 1 | JAB75945 | Ribosomal protein LP2 [*Ixodes ricinus*] |
| ORTHOMCL3820 | 3 | 1 | 1 | ISCW002028 | IMP-GMP specific 5'-nucleotidase |
| ORTHOMCL4006 | 1 | 1 | 3 | EEC13628 | Urocanate hydratase |
| ORTHOMCL3946 | 1 | 1 | 3 | EEC20451 | Uncharacterised protein (pseudouridine-5'-phosphate glycosidase domain) |
| ORTHOMCL5111 | 2 | 1 | 1 | ISCW010907 | Formiminotransferase-cyclodeaminase |
| ORTHOMCL75 | 27 | 1 | 0 | EEC02489 | Cysteine-rich secreted protein (trypsin inhibitor-like cysteine rich domain) |
| ORTHOMCL8850 | 2 | 1 | 0 | ISCW012704 | Signal sequence receptor beta |

<sup>a</sup>Annotations are from *I. scapularis* unless otherwise stated. Protein domain information was obtained from the Conserved Domain Database [228]. Tick salivary protein data were obtained from Kim *et al.* [120].

Several other protein clusters with potential roles in immune evasion or the regulation of salivation and the ingestion of host fluids were identified in the trombidid mite genomes. The presence of an expanded acetylcholinesterase gene family in tick genomes has been noted previously and acetylcholinesterases have been detected in the saliva of *Rhipicephalus microplus* [129] and *Amblyomma americanum* [122], as well as *I. scapularis* [120]. It has been proposed that salivary acetylcholinesterases could interfere with cholinergic signalling between host immune cells and might facilitate pathogen establishment [129]. However, the trombidid mites have only a single gene copy each that clusters with the *I. scapularis* acetylcholinesterases (Table 3). Similarly, ATPase inhibitors (Table 3) in saliva could impact on local immune responses [122], since extracellular purine metabolites act as “alarmins” [130]. The massive expansion of sulfotransferases in the *I. scapularis* genome and the secretion of some members of this family in saliva is particularly enigmatic, but recently it has been proposed that sulfotransferases could control salivation and feeding cycles in ticks by sulphating the neurotransmitters dopamine and octopamine [131]. Alternatively or in addition, they might be involved in increasing the activity of small cysteine-free thrombin inhibitors in tick saliva by sulfation of tyrosine residues [132]. Notably, only two of these sulfotransferases were present in each trombidid mite genome compared with >70 members of this family in *I. scapularis* (Table 3).

Of the two salivary protein families restricted to *L. deliense* and *I. scapularis* (Table 3), the secreted trypsin inhibitor-like cysteine-rich domain proteins are among are wide diversity of serine protease inhibitors produced by ticks [133]. This specific class of trypsin inhibitor-like proteins includes ixodidin, an antimicrobial peptide expressed in the haemocytes of *R. microplus* [134] and BmSI-7 and BmSI-6, two peptides from the same species of tick that inhibit cuticle-penetrating proteases secreted by entomopathogenic fungi [135]. The BmSI-7 peptide is expressed in multiple tissues, including the salivary glands [135], but its role in saliva is unknown. However, one possibility is that it helps prevent the tick bite site from becoming infected. In contrast with *I. scapularis*, which harbours 27 trypsin inhibitor-like proteins in its genome, only one orthologue was identified in *L. deliense* (Table 3). The second cluster restricted to *I. scapularis* and *L. deliense*, a signal sequence receptor subunit (Table 3), was unexpected as it has a canonical function in trafficking secretory proteins through the ER [136]. This appears to have a moonlighting role in tick saliva, since it generates strong immune responses in rabbits parasitized by *A. americanum* [122].

Feeding ticks secure their mouthparts in the skin of the host for days or weeks using a cement-like substance that forms a cone in the bite wound. Superficially, the stylostome generated at the feeding site of trombidid mites resembles the tick cement cone, although the structure is tubular (in the Trombiculoidea) or highly branched (in the Trombidioidea) [40]. In both of the trombidid mite genomes, we found an orthologue of a glycine-rich protein present in the sialome of *Rhipicephalus pulchellus* [124] (Table 3). The tick glycine-rich proteins are related to spider silk proteins and form the main structural component of tick cement as determined by proteomic studies [137]. To determine if the trombidid mite genomes may contain other cement-associated proteins not detected in the *I. scapularis* salivary proteomics study [120], we searched for orthologues of all tick cement proteins in the National Center for Biotechnology Information (NCBI) database. We found orthologues of an *I. scapularis* glycine-rich cement protein in both *L. deliense* (one copy) and *D. tinctorium* (three copies) that was distinct from the *R. pulchellus* orthologue; moreover, the velvet mite also possessed an orthologue of a second *I. scapularis* cement protein (Additional file 20). In addition, both trombidid mites harboured a gene related to a cement protein transcript identified in the sialotranscriptome of *Amblyomma triste* [125] (Additional file 20). Finally, orthologues of *A. americanum* acidic chitinases involved in conferring stability to the tick cement cone were present in both *D. tinctorium* (four copies) and *L. deliense* (one copy) [138] (Additional file 20).

## Discussion

### Genome features and trombidid mite evolution

In this study, we exploited the close phylogenetic relationship between the Trombidioidea and the Trombiculoidea in order to obtain a genome from a single adult velvet mite that could be used to corroborate data derived from a suboptimal trombiculid mite sample (i.e., a pool of engorged larvae). This strategy proved successful because in almost all cases, the unusual features of the trombidid mite genomes were shared between the two sequenced taxa. In contrast with other acariform mites, the trombidid mite genomes were substantially larger, contained a greater proportion of repeats, and exhibited expansions of mobile elements. These features, coupled with heterozygosity and host contamination in the case of *L. deliense*, proved challenging for accurate genome size estimation but did not prevent the annotation of protein-coding genes, which was sufficient (even for *L. deliense*) for an initial protein expression study. Our *k*-mer-based estimate of genome size for *L. deliense* was slightly smaller than those determined for *Leptotrombidium pallidum* and *Leptotrombidium scutellare* using DNA from laboratory-reared adult specimens, which were 191 ± 7 Mb and 262 ± 13 Mb (by qPCR), or 175 Mb and 286 Mb (by *k*-mer analysis), respectively [139]. In future, genome sequencing from individual *L. deliense* adults would help resolve the variation in genome size across the *Leptotrombidium* genus, which is likely to be driven by repeat content.

As previously reported on publication of the first spider genomes [61], the Acari are polyphyletic, with the superorder Parasitiformes (*i.e.*, the ticks together with mesostigmatid and holothyrid mites) evidently more closely aligned to the spiders (order Araneae) than it is to the Acariformes (Figure S1). However, at lower taxonomic scales, the phylogenomic analyses supported the conventional morphology-based taxonomy within the Acariformes, confirming that the trombidid mites are closely related to the phytophagous Tetranychoidea ([14]. Interestingly, within the Trombidiae, the divergence between the velvet mites and the chiggers occurred much later (~123 MYA) than the emergence of the earliest terrestrial vertebrates (395 MYA [140]), coinciding perhaps with the appearance of crown-group mammals. If this scenario is correct, the ectoparasitism of non-mammalian vertebrates by chiggers that occurs today may be a product of secondary adaptation, as suggested by other authors [20]. Although the fossil record is devoid of trombidid mite specimens predating the Eocene [141], it has been speculated from palaeogeographical and comparative morphological evidence that trombiculid mites fed initially on other arthropods, with larval ectoparasitism on vertebrates evolving during the Paleocene, leading to an increase in chigger diversity [20]. Our data challenges this hypothesis, because it implies that something other than host choice in the larval stage drove the split between the Trombidioidea and the Trombiculoidea 60 million years before the latter began feeding on vertebrates. Only the discovery of more ancient trombidid fossils will help to resolve these uncertainties.

### Potential roles for terpenes in trombidid mite biology

The most striking finding in the trombidid mite genomes was the presence of large families of laterally-transferred terpene synthases. As the level of amino acid identity between the trombidid terpene synthases and their closest homologues in microbes is quite low, their end products cannot be inferred with any confidence. However, the question of whether compounds such as 2-methylisoborneol, geosmin or germacrene might confer adaptive advantages to trombidid mites helps to frame hypotheses for experimental testing. Interestingly, all of these compounds are associated with odours and tastes that humans, and some arthropods, may sense as unpleasant or aversive. For instance, 2-methylisoborneol has a musty odour that humans associate with ripe cheeses [142]; whereas geosmin confers the smell of moist soil and is the cause of the muddy, “off” taste that the human olfactory system detects in spoiled water, wine and the flesh of certain freshwater fish [143-145]. More importantly, geosmin released by *Penicillium* spp. or *Streptomyces* spp. on rotting fruit is strongly aversive to *Drosophila* because these organisms produce secondary metabolites that are directly toxic to the fly or to its primary food source, yeast [146]. Germacrene has also been implicated as an arthropod repellent amongst complex sesquiterpene mixtures found in the essential oils of various plants, which have been shown to be effective against several acarines, including the ticks *Rhipicephalus microplus* and *Ixodes ricinus* [147, 148], and the poultry red mite *Dermanyssus gallinae* [148]. Although it has not been assayed in isolation, germacrene is additionally a significant component of essential oils or crude leaf extracts that exhibit toxic effects against phytophagous mites (*Brevipalpus phoenicis* in the Trombidiformes [149]) and ants (*Solenopsis invicta* [150]).

These potential repelling and/or toxic effects of terpenoids would align closely with the apparent aposomatic nature of trombidid mites, most species of which are brightly coloured due to their carotenoid content. These mites have few natural enemies and have been reported to be rapidly regurgitated if offered to predators in the laboratory [151]. However, cannibalism between adults and even ectoparasitism of the free-living stages by trombidid larvae can occur, underlining the relevance of chemical communication within species and between closely related species [18]. To the best of our knowledge, only one report of a parasitoid affecting trombidid mites has been published (the acrocerid fly *Pterodontia flavipes* attacking *Podothrombium* spp. [152]), but the dearth of research on free-living trombidid stages means that no doubt other parasitoids exploiting these hosts do exist. It is important to note that not all mites are repelled by terpenoid compounds. Many plants use terpenoids as defence compounds to signal to the natural enemies of pest arthropods that the plant is under attack. For example, *Lotus japonicus* infested with *T. urticae* releases several terpenoids, including germacrene, which attract the predatory mite *Phytoseiulus persimilis* [153]. Communication by sex pheromones is also known to occur in *T. urticae*, although molecules other than terpenoids are suspected to mediate this [154]. Moreover, as noted above, neryl formate is an aggregation pheromone in dust mites [76]. In conclusion, while the conferment of a foul taste (and perhaps odour) to the aposomatic trombidids appears to be the most likely evolutionary driver of *de novo* terpenoid synthesis capability, it is also possible that these compounds are used to communicate with potential mates during courtship, or to repel members of the same (or closely-related) species to reduce competition or deter cannibalistic behaviour.

### Diapause and seasonality

Many of the putative functions of laterally transferred and/or expanding gene families in the trombidid mites appear to be relevant to the temporal regulation of the lifecycle and the switching of metabolic demands between dormant and active stages. The life history of trombidid mites features alternation between immobile calyptostases (the deutovum, protonymph and tritonymph) and the active instars (larva, deutonymph and adult) [155]. The calyptostases typically persist for 25 – 30 days, while the active stages in temperate species can undergo hibernation over the winter months, including larvae that have not fed until late in the autumn. Diapause of eggs is common in temperate species and can exceed one year in the chigger, *Hirsutiella zachvatkini*, without loss of viability [20]; while larvae of this species can also overwinter on their rodent hosts [156]. The lifecycle of trombidid mites in tropical and subtropical regions has been little studied, but several generations per year are possible [20]. In the case of *Dinothrombium* spp., although the adults are positively phototactic and diurnal if humidity is high (becoming crepuscular during drier conditions), circadian cycles of activity were maintained if mites were transferred to constant darkness in the laboratory [151]. Remarkably, adult *D. pandorae* of the Californian deserts may only emerge from their burrows during rainstorms to feed and mate for a few hours each year, migrating to the deepest extent of their subterranean refuges during the height of summer [43]. However, overcoming torpor by rapidly adjusting metabolic rate after the cold desert night to the early morning warmth, when termite prey become active, is critical to the lifecycle of *D. pandorae* [43]. It may also be important for chiggers to avoid cool microclimates (and thus maintain peak metabolism) when questing for small mammals, since their hosts are highly motile and a suitable location for attachment must be targeted rapidly before grooming behaviour leads to ingestion of the mite. Hence, the small expansions in the IR repertoire (IR25a, IR21a and IR93a) that we observed in the trombidid mite genomes might reflect more acute sensitivity to cool temperatures than for the phytophagous *T. urticae*.

With relevance to both regulation of metabolism and circadian cycles, the homologues of FLVCR proteins have been little studied in arthropods, but in vertebrates they export haem from the cytoplasm to the extracellular milieu (for FLVCR1) and from mitochondria into the cytoplasm (for FLVCR2) [157]. A FLVCR gene homologue in *Drosophila melanogaster, CG1358*, is involved in maintenance of circadian rhythms in the absence of light together with other genes with roles in iron metabolism [158]. Thus, it is intriguing that FLVCR homologues were significantly expanded in the *D. tinctorium* genome but not that of *L. deliense*, as adult chiggers in tropical environments exhibit more regular activity above ground than do *Dinothrombium* spp. [20]. To the best of our knowledge, the function of the 4-coumarate:CoA ligase family has not been explored in the Arachnida, but in adult females of the kissing bug *Rhodnius prolixus*, RNAi of long-chain acyl-CoA synthetase 2 led to a 90% decrease in fatty acid β-oxidation and substantial reductions in oviposition and egg hatching, as well as marked abnormalities in the remaining eggs and hatched nymphs [159]. The large gene expansions in acyl-CoA synthetases observed here in *D. tinctorium* suggest that β-oxidation is a particularly important facet of its metabolism. Furthermore, the proteomic analysis of *D. tinctorium* revealed overrepresentation of putative digestive enzymes with peptidase M20 and inhibitor I29 domains, which is consistent with an imperative for adult velvet mites to obtain food reserves rapidly while foraging briefly above ground [43]. An elevated metabolic rate for *L. deliense* larvae was also suggested by the preponderance of mitochondrial enzymes responsible for aerobic energy production in protein extracts. Further studies on trombidid mite metabolism are clearly warranted, as most metabolic studies in the Trombidiformes have focused on winter diapause in *T. urticae* [160], creating a knowledge gap around the physiology of tropical species.

In spider mites, carotenoids are essential for the control of diapause and sexual reproduction. However, reproductive behaviour between the Tetranychoidea and the trombidid mites is radically different, since male *T. urticae* become developmentally arrested in close proximity to dormant female deutonymphs on leaf surfaces. This “guarding” behaviour is stimulated by the intensity of the yellow colouration of the dormant females (derived from their carotenoid pigments), and allows the male to mate with the adult female immediately after ecdysis [161]. In contrast, in order to be inseminated, adult females of trombidid mites must collect a spermatophore deposited on the ground by the male, the location of which is signposted by signalling threads. While colouration is not known to be factor during courtship, the males of some species deposit spermatophores in specially constructed “gardens” and perform encircling dances with the female [18]. However, it has recently been discovered using genetic manipulation of a laterally-transferred phytoene desaturase that carotenoids have a second, distinct function in the regulation of diapause in *T. urticae* [13]. A lack of phytoene desaturase activity not only results in albinism, but prevents overwintering strains from entering diapause, probably due to disrupted light (and thus photoperiod) perception caused by vitamin A deficiency.

To the best of our knowledge, anatomical and experimental studies on trombidid mite vision have not been performed, but spider mites once again provide a closely-related template. The eyes of *T. urticae* have been partially characterised, and show biconvex lenses in the anterior pair and simplex convex lenses in the posterior pair [162]. It has been proposed that the anterior eyes respond to UV and green light, whereas the posterior pair are sensitive to UV only. However, laser ablation experiments have demonstrated that either pair can receive sufficient information to control the photoperiodic termination of diapause, while removal of both pairs prevents diapausal exit [163]. In *T. urticae*, the expression pattern of rhodopins in the eyes has not been determined, but in the jumping spider *Hasarius adansoni*, green-sensitive rhodopsin-1 expressed in the lateral ocelli is important for the monochromatic detection of movement [164]. Until recently, the function of rhodopsin-7 was enigmatic, but experiments in *D. melanogaster* have shown that it operates in the circadian pacemaker neurons of the central brain and is responsible for their highly sensitive response to violet light [165]. Taken together, these experimental findings from other arthropods suggest that trombidid mites might depend on rhodopsin-7 homologues rather than peropsins for control of circadian rhythms (although we failed to identify orthologues of the *Drosophila* clock gene in Acari). The relative roles of FLVCR gene homologues, phytoene desaturases and the rhodopsin pigments in the control of diapause and other life history traits in trombidid mites is evidently a key priority for future experimental studies.

### Immune response and vector biology

As trombidid mites are edaphic organisms with a parasitic larval stage, they are exposed to soil microorganisms, the exterior flora of their hosts, and pathogens contained in ingested body fluids. In the case of certain trombiculid mites (especially *Leptotrombidium* spp.), their role as biological vectors of *O. tsutsugamushi* highlights a specific infectious challenge that has resulted from feeding on small mammals. In other arthropods, CTLD proteins with lectin activity act as transmembrane or secreted pattern recognition receptors that bind to carbohydrates on the surface of pathogens. They can function as opsonins in the haemolymph that agglutinate unicellular pathogens and facilitate their phagocytosis by haemocytes [166], or may be expressed on the surface of tissues that form a barrier to infectious assaults, such as in the gut or on the gills of crustaceans [92, 167]. These findings are compatible with roles as secreted opsonins for most of the CTLD proteins identified in the current study, with a smaller number perhaps operating as immune surveillance receptors on the surface of cells or extracellular matrices.

It is unclear why the *L. deliense* genome harbours a variety of PERK-like proteins. However, the alpha subunit of eukaryotic translation-initiation factor-2 is the main substrate of PERK and is inactivated by phosphorylation, leading to inhibition of global protein synthesis. As the unfolded protein response bridges the cellular response to viral infection and metabolic homeostasis [168], the manipulation of PERK by dengue virus [169] suggests that an expanded repertoire of these kinases might be beneficial in the evolutionary arms race with viral pathogens. Indeed, we identified a striking diversity of ERV-related sequences in the *L. deliense* genome, apparently reflecting a substantial degree of exposure to viral nucleic acids. The relatively large number of these Gag-like polyproteins, the absence of similar quantities of other retroviral proteins in the *L. deliense* assembly, and the closer relationship of the Gag-like polyproteins to ERV elements in non-sciurid mammals, all rendered host DNA contamination in the mite gut or on mouthparts as an unlikely source for these sequences; although this possibility cannot be excluded entirely. Despite this caveat, lateral transfers in the distant past originating from mammalian body fluids during the brief ectoparasitic stage is a working hypothesis that can be tested when a more contiguous chigger genome becomes available. In contrast, the horizontal transfer of the long-interspersed element BovB is postulated to have involved an opposite transmission route; that is, between vertebrates by ticks [170].

To the best of our knowledge, no experimental studies on the immune response of trombidid mites have been performed to date. However, a recent study in *T. urticae* involving experimental challenge with bacteria demonstrated that the spider mites were highly susceptible to systemic infection [171]. This contrasted with a more robust response to bacterial challenge in another acariform mite, *Sancassania berlesei*, which unlike *T. urticae* has a saprophytic lifestyle. The authors of this study concluded that the ecology of spider mites, in which all lifecycle stages feed on plant phloem (a relatively aseptic food source), has led to a high degree of susceptibility to pathogen exposure. This failure to overcome infectious insults was associated with an apparent absence of many antimicrobial protein effectors in the spider mite genome. In support of the hypothesis that spider mites have adapted to an environment characterised by very low levels of pathogen challenge, we found that compared with the *L. deliense* genome, the *T. urticae* genome displays a relative paucity of PGRP (Additional file 17), CTLD (Additional file 6) and Dscam genes (Additional file 16). Thus, the common ancestor of the Trombidiformes may have harboured a diverse immune gene repertoire that was selectively lost in the branch leading to the Tetranychoidea, and/or the Trombiculoidea has undergone more recent immune gene family expansions. The intermediate immune gene repertoire of the Trombidioidea between that of the spider mites and the chiggers (*D. tinctorium* has considerably fewer CTLD proteins and PERKs than *L. deliense*) suggests that both immune-related gene losses and gains have occurred during the evolution of the Trombidiformes in response to their radically different natural histories. Indeed, in terms of the degree of exposure to pathogen diversity and abundance, the euedaphic velvet mites are likely to encounter greater infectious challenges than spider mites, and the feeding behaviour of chiggers on vertebrates is likely to exacerbate this exposure further compared to their relatives that are ectoparasitic on other invertebrates only.

## Conclusions

This first analysis of trombidid mite genomes has revealed their dynamic nature relative to those of other acariform mites, including expansions in laterally-transferred gene families and mobile elements. These genomes provide a foundation for fundamental experimental studies on mite immune responses, host-seeking behaviour and feeding, and environmental impacts on lifecycle progression. The function of the laterally-transferred terpene synthases will become a major research theme for chigger biology, as only experimental exposure of chiggers and their potential natural enemies to mite terpene extracts will be able to determine if these unique aspects of secondary metabolism have evolved to attract conspecifics; or conversely, to repel predators, parasitoids and/or competitors. From an applied perspective, the identification of predicted allergens in the *L. deliense* genome sets the scene for immunoproteomic studies of trombiculiasis in both humans and domestic animals, with the potential for immunotherapeutic approaches to be developed as for dust mite allergy [172]. Finally, the successful development of recombinant vaccines against ticks [173] and the promising progress of recombinant vaccine development for both sheep scab [174] and poultry red mite [175] indicate that a similar approach could be explored for chiggers, which has the potential to interrupt, or at least reduce, the transmission of *O. tsutsugamushi* to humans in scrub typhus-endemic areas. Considering the high strain variability of the scrub typhus agent [176], a chigger vaccine utilising mite salivary or gut antigens could provide a much-needed breakthrough against this intractable disease.

## Materials and methods

### Sample collection and DNA extraction

Adult specimens of giant red velvet mites were collected within the grounds of the UK Medical Research Council Field Station at Wali Kunda, The Gambia (13°34’N, 14°55’W). Mites were sampled from flowerbeds following heavy rains in June 2010 and stored in 95% ethanol at −80°C. They were identified as *Dinothrombium tinctorium* by Joanna Mąkol (Wrocław University of Environmental and Life Sciences, Poland). Approximately 5 µg of DNA was extracted from a single individual using a Genomic-tip Kit (Qiagen) according to the manufacturer’s instructions. Integrity of the DNA was confirmed by agarose gel electrophoresis, which showed a single band of ~20 kb.

For *L. deliense*, engorged larvae were collected from two Berdmore’s ground squirrels (*Menetes berdmorei*) captured in Udonthani Province, Thailand, in September 2015. Trapping and euthanasia of small mammals followed the CERoPath (Community Ecology of Rodents and their Pathogens in a changing environment) project protocols [177]. Chiggers were located inside the ears and the inguinal area of the squirrels and stored in absolute ethanol at −20°C. A subsample of the mites was selected and mounted in clearing medium, Berlese fluid (TCS Bioscience, UK), prior to species identification under a compound microscope. Fifty unmounted larvae were pooled and ~30 ng of genomic DNA was extracted using a DNeasy Blood & Tissue Kit (Qiagen) according to the manufacturer’s instructions. The DNA was partially degraded, but a dominant band of ~5 kb was apparent by agarose gel electrophoresis.

### Library preparation and sequencing

These steps were performed at the Centre for Genomic Research at the University of Liverpool. The *D. tinctorium* DNA was used to generate two Illumina TruSeq libraries and one Nextera mate-pair library. For the former, bead-based size selection using 100 ng and 200 ng of DNA as input into the TruSeq DNA LT Sample Prep Kit with 350 and 550 bp inserts, respectively, was applied. Following eight cycles of amplification, libraries were purified using Agencourt AMPure XP beads (Beckman Coulter). Each library was quantified using a Qubit fluorimeter (Life Technologies) and the size distribution was assessed using a 2100 Bioanalyzer (Agilent). The final libraries were pooled in equimolar amounts using the Qubit and Bioanalyzer data. The quantity and quality of each pool was assessed on the Bioanalyzer and subsequently by qPCR using the Illumina Library Quantification Kit (KAPA Biosystems) on a LightCycler 480 instrument II (Roche Molecular Diagnostics) according to the manufacturer’s instructions. The pool of libraries was sequenced on one lane of the HiSeq 2000 with 2 × 100 bp PE sequencing and v3 chemistry.

The *D. tinctorium* mate-pair library was constructed using the Nextera Mate Pair Kit (Illumina) with 3 kb inserts. The DNA (3 µg) was tagmented as described in the manufacturer’s protocol and cleaned using a Genomic DNA Clean & Concentrator column (Zymo Research). The sample was then subjected to strand displacement and cleaned with AMPure XP beads. A 0.6% Certified Megabase Agarose gel (Bio-Rad) was used to separate the fragments, and those in the range of 2 – 5 kb were extracted and recovered using a Zymoclean Large Fragment DNA Recovery Kit (Zymo Research). The recovered DNA was quantified and transferred into a circularisation reaction at 16°C overnight. After purification with AMPure XP beads, DNA was sonicated into ~500 bp fragments using a focused ultrasonicator (Covaris) and recovered with AMPure XP beads as before. Samples were bound to Dynabeads M-280 Streptavidin (Thermo Fisher Scientific) and all subsequent reactions (end repair, A-tailing, and adapter ligation) were bead-based. Samples were amplified with 10 cycles of PCR, recovered by AMPure XP beads at a 1:1 ratio, and quantified using the Qubit dsDNA HS Assay Kit. The library was then subjected to quality control on the Bioanalyzer and LightCycler as for the TruSeq libraries above. The library was sequenced on one run of the MiSeq with 2 × 250 bp PE sequencing.

For *L. deliense*, the DNA sample was sheared to 550 bp using a Bioruptor Pico sonication device (Diagenode) and purified using an AxyPrep FragmentSelect-I Kit (Axygen). The sample was then quantified using a Qubit dsDNA HS Assay Kit on the Qubit fluorimeter, and the size distribution was ascertained on the Bioanalyzer using a High Sensitivity DNA chip (Agilent). The entire sample was used as input material for the NEB Next Ultra DNA Library Preparation Kit. Following nine PCR cycles, the library was purified using the AxyPrep kit and quantified as before by Qubit. Library size was determined on the Bioanalyzer (Agilent). The quality and quantity of the pool was assessed as described above for *D. tinctorium*. The sequencing was conducted on one lane of an Illumina MiSeq with 2 × 150 bp PE sequencing and v2 chemistry.

### Assembly and annotation

For both genomes, base-calling and de-multiplexing of indexed reads was performed by bcl2fastq v. 1.8.4 (Illumina) to produce sequence data in fastq format. The raw fastq files were trimmed to remove Illumina adapter sequences using Cutadapt version 1.2.1 [178]. The option “-O 3” was set, so the 3’ end of any reads which matched the adapter sequence over at least 3 bp was trimmed off. The reads were further trimmed to remove low quality bases, using Sickle version 1.200 [179] with a minimum window quality score of 20. After trimming, reads shorter than 10 bp were removed. If both reads from a pair passed this filter, each was included in the R1 (forward reads) or R2 (reverse reads) file. If only one of a read pair passed this filter, it was included in the R0 (unpaired reads) file.

The Kraken taxonomic sequence classification system (v. 1.0) [180] was used to assign taxonomic annotations to the reads and specifically to estimate the level of bacterial contamination in the raw data. For genome size estimations, *k*-mers were counted by the Jellyfish program (v. 2.0) [181] and the resultant histograms were uploaded to GenomeScope (v. 1.0) [182] to visualise the *k*-mer distributions.

For *D. tinctorium*, the PE reads were assembled using Abyss (v. 1.5.2) [48, 49], Allpaths-LG (v. r51279) [183], SOAPdenovo2 (v. 2.04-r240) [184] and Discovar (v. r51454) [185]. When running Abyss, *k*-mer sizes were from 35 bp to 80 bp with an interval of 5, and the output “*k*-mer size = 80 bp” was selected as this produced the optimal assembly. Allpaths-LG and Discovar specify *k*-mer size automatically, whereas “*k*-mer size = 63” was selected for SOAPdenovo2, as suggested by the developer. Discovar requires read lengths of 250 bp, so was applied only to the data generated from the mate-pair library. Assessment of the completeness of the genome assemblies was based on the percentage alignment obtained against the reads from the TruSeq 350-bp insert library using bowtie2 (v. 2.0.10) [186] and the predicted core gene content determined by CEGMA (Core Eukaryotic Genes Mapping Approach) (v. 2.5) [187]. The final, optimum assembly was created by Abyss (97.4% of scaffolds >500 bp were mapped; 99.2% of Key Orthologs for eukaryotic Genomes were present) and included all scaffolds of ≥1,000 bp, which constituted ~80% of the total length of the raw assembly.

For *L. deliense*, a preliminary genome assembly at contig level was performed using Velvet (v. 1.2.07) [50], with parameters of ‘best *k*-mer 99’ and ‘-ins_length 500’. Reads derived from mammalian host genomic DNA were removed from the preliminary genome assembly using blobtools (v0.9.19), which generates a GC-coverage plot (proportion of GC bases and node coverage) [51] (Fig. 3). The filtered *L.deliense* reads were then reassembled using SPAdes assembler (v3.7.1) [52] with default settings and the length cut-off for scaffolds was set at 500 bp.

To find, classify and mask repeated sequences in the mite genome assemblies, a *de novo* repeat library was first built using RepeatModeler (v. 1.0.8) [188] with the ‘-database’ function, followed by application of RepeatMasker (v. 4.0.6) [189] using default settings for *de novo* repeated sequences prediction. Then, a homology-based prediction of repeated sequences in the genome was achieved using RepeatMasker with default settings to search against the RepBase repeat library (issued on August 07, 2015). For non-interspersed repeated sequences, RepeatMasker was run with the ‘-noint’ option, which is specific for simple repeats, microsatellites, and low-complexity repeats.

Thee *ab initio* gene prediction programs, including Augustus (v. 3.2.2) [54], SNAP (v. 2013-11-29) [55] and GeneMark (v. 2.3e) [56] were used for *de novo* gene predictions in each genome assembly. Augustus and SNAP were trained based on the gene structures generated by CEGMA (v. 2.5) [187], whereas GeneMark [56] was self-trained with the ‘–BP OFF’ option. All three *ab initio* gene prediction programs were run with default settings. We also generated an integrated gene set for each genome assembly using the MAKER v. 2.31.8 [53] pipeline. The MAKER pipeline runs Augustus, SNAP and GeneMark to produce *de novo* gene predictions, and integrates them with evidence-based predictions. These were generated by aligning the invertebrate RefSeq protein sequences (downloaded on March 31, 2016 from NCBI) to the masked mite genomes by BLASTX. The MAKER pipeline was run with ‘-RM_off’ option to turn all repeat masking options off, and all parameters in control files were left in their default settings. Genes identified by *de novo* prediction, which did not overlap with any genes in the integrated gene sets, were also added to the final gene set for each genome assembly if they could be annotated by InterProScan (v. 4.8) [190] with the InterPro superfamily database (v. 43.1) using ‘-appl superfamily -nocrc’ options.

The Blast2GO pipeline (v. 2.5) [191] was used to annotate proteins by Gene Ontology (GO) terms. In the first step, all protein sequences were searched against the nr database with BLASTP. The E-value cutoff was set at 1 × 10^−6^ and the best 20 hits were used for annotation. Based on the BLAST results, the Blast2GO pipeline then predicted the functions of the sequences and assigned GO terms to the BLAST-based annotations. Metabolic pathways were constructed using KAAS (KEGG Automatic Annotation Server) [192] with the recommended eukaryote sets, all other available insects, and *I. scapularis*. The pathways in which each gene product might be involved were derived from the best KO hit with the BBH (bi-directional best hit) method.

### Phylogenetics

Protein data sets of the following arthropod genomes were used as references: *D. melanogaster* (fruit fly; GOS release: 6.11) [193], *A. mellifera* (honey bee; GOS release: 3.2) [194], *T. mercedesae* (bee mite; GOS release: v. 1.0) [98], *T. urticae* (spider mite; GOS release: 20150904) [59], *Stegodyphus mimosarum* (velvet spider; GOS release: 1.0) [61], *I. scapularis* (blacklegged tick; GOS release: 1.4) [195], *M. occidentalis* (predatory mite; GOS release: 1.0) [97]. *Caenorhabditis elegans* (nematode; GOS release: WS239) [196] was used as the outgroup. For gene-family phylogenetics, we first aligned orthologous protein sequences with Mafft (v7.309) [197] or Kalign (v2.0) [198]. We manually trimmed the aligned sequences for large gene sets. The best substitution models of amino-acid substitution were determined for the alignments by ProtTest (v3.4) with parameters set to “-all-matrices, -all-distributions, -AIC” [199]. Then, phylogenetic trees were constructed using maximum likelihood methods (Phyml, v3.1) [200]. In addition, a neighbour-joining method was used for building the distance-based trees using MEGA (7.021) [201].

For species-level phylogenetics, the rapid evolution of acariform mites may challenge phylogenetic analyses due to long-branch attraction [202]. Thus, we used a very strict E-value (1 × 10^−50^) when performing a reciprocal BLASTP to exclude the most variant orthologous genes across all genomes tested. The reciprocal BLAST search resulted in identification of a total of 926 highly conserved one-to-one orthologues in all eight genomes. Each of these orthologous groups was aligned using Mafft with the “-auto” option. These alignments were trimmed by Gblocks (v. 0.91b) [203] and concatenated into unique protein super-alignments. ProtTest determined the best-fit substitution model of LG with invariant sites (0.131) and gamma distributed rates (0.913) using parameters as above prior to conducting the phylogenetic analysis with Phyml and Bayesian methods (MrBayes, v. 3.2.6) [204]. Based on the topology defined by this phylogenetic analysis, we estimated the divergence time of each species using the Bayesian MCMC method in the PAML package (v. 4.9a) [205] with the correction of several fossil records (time expressed in MYA): tick-spider: 311–503 [61] (oldest spider from coal, UK), *T. urticae*-tick-spider: 395–503 [61] (oldest Acari), *A. mellifera-D. melanogaster:* 238-307 [206] and nematode-arthropods: 521–581 [206].

### Analysis of gene family expansions

Orthologous gene clusters of *D. tinctorium, L. deliense* and the other reference genomes described above were defined based on OrthoMCL (v. 1.4) [207]. We used CAFE (v. 3.1) [208] to infer the gene family expansion and contraction in *D. tinctorium* and *L. deliense* against other Acariformes (*T. urticae* and *S. scabiei*). The ultrametric species tree used in the CAFE analyses was created as described for gene-family phylogenetics above. We also calculated ω (*d*_N_/*d*_S_) ratios for 454 one-to-one orthologues defined by OrthoMCL using codeml in the PAML package [205] with the free-ratio model. Branches with ω >1 are considered to be under positive selection. The null model used for the branch test was the one-ratio model (nssites = 0; model = 0) where ω was the same for all branches. Kappa and omega values were automatically estimated from the data, with the clock entirely free to change among branches. The *P* value was determined twice using the log-likelihood difference between the two models, compared to a χ^2^ distribution with the difference in number of parameters between the one-ratio and free-ratio models. To estimate significance with the *P* value, a likelihood-ratio test was used to compare lnl values for each model and test if they were significantly different. The differences in log-likelihood values between two models were compared to a χ^2^ distribution with degrees of freedom equal to the difference in the number of parameters for the two models. Measurement of *d*_s_ was assessed for substitution saturation, and only *d_s_* values <3.0 were maintained in the analysis for positive selection. Genes with high ω (>10) were also discarded.

### Analysis of candidate lateral gene transfers

We used a modification of the Crisp method [6] for examination of LGTs in the two mite genomes. Each mite protein dataset was aligned with BLASTP against two databases derived from the NCBI nr database, one consisting of metazoan proteins (excluding proteins from species in the same phylum as the studied species - Arthropoda) and the other of non-metazoan proteins. The LGT index, h, was calculated by subtracting the bit-score of the best metazoan match from that of the best non-metazoan match. The genes can be classified into class C if they gained an *h* index ≥ 30 and a best non-metazoan bit-score of ≥100. For each class C gene, its average *h* value (h_orth_) and that of its paralogous genes in each OrthoMCL cluster defined above was determined. If *h* was ≥30, the best non-metazoan bit-score was ≥100, and the h_orth_ value was ≥30, the gene was considered to be a class B gene. Class A genes were defined as a subset of class B genes with *h* ≥30, a best non-metazoan bit-score of ≥100, an h_orth_ value of ≥30, and a best metazoan bit-score of <100.

### Analysis of immune-related gene families

A search for mite immune-related genes was initially preformed with a BLASTP search (E-value, <1 × 10^−5^) against each mite protein set using immune-related genes defined by Palmer & Jiggins [85]. The identified potentially immune-related genes were then manually checked using BLASTP online at NCBI. For analysis of CTLD proteins, FASTA sequences of proteins in ORTHOMCL136, ORTHOMCL510, ORTHOMCL582, and ORTHOMCL880 were analysed by SecretomeP 2.0 [209], TMHMM 2.0 [210] and NetNGlyc 1.0 [211] using default settings to identify respective sequence features. Protein sequences were also submitted to InterPro [212] for domain structure analysis. The CTLDs extracted from individual protein sequences were then manually searched for the amino-acid motifs “EPN”, “WND” and “QPD”.

### Analysis of chemosensory and photoreceptor gene families

A search for *D. tinctorium* and *L. deliense* OBPs was initially preformed using TBLASTN (E-value, <1 × 10^−3^) against their genome assemblies using *D. melanogaster, Drosophila mojavensis, Anopheles gambiae, Bombyx mori, Tribolium castaneum, A. mellifera, Pediculus humanus humanus* and *Acyrthosiphon pisum* OBPs (identified by Vieira and Rozas [213]) as queries. No OBPs were found in the *D. tinctorium* and *L. deliense* genome assemblies. Because OBPs are very divergent in terms of the amino-acid sequences within the family, and the sequence identities between the family members from the different species can be as low as 8% [213], a TBLASTN search has limited power to identify these genes. A search for OBPs was therefore performed again with BLASTP (E-value, <1 × 10^−3^) to search the automated protein predictions from the mite genome assemblies. A search for small chemosensory proteins is *D. tinctorium* and *L. deliense* was preformed using the same methods as for the OBPs. The query sequences were also based on the study of Vieira and Rozas [213], using chemosensory protein sequences from *D. melanogaster, D. mojavensis, A. gambiae, B. mori, T. castaneum, A. mellifera, P. humanus humanus, A. pisum, I. scapularis*, and *Daphnia pulex*. For odorant receptors, both TBLASTN and BLASTP searches were performed using *D. melanogaster* and *A. mellifera* sequences (identified by Nozawa and Nei [214] and Robertson and Wanner [215], respectively) as queries.

The *D. tinctorium* and *L. deliense* GR gene families were manually annotated according to TBLASTN and BLASTP searches (both with an E-value cutoff of <1 × 10^−3^) against their genome assemblies and predicted protein coding genes, respectively, using all *D. melanogaster* [216], *A. mellifera* [215], *I. scapularis* [195], *T. urticae* [100], *T. mercedesae* [98] and *M. occidentalis* [97] GRs as queries. An iterative search was also conducted with termite GRs as queries until no new genes were identified in each major subfamily or lineage. For phylogenetic analysis, *D. tinctorium* and *L. deliense* GRs were aligned with *D. melanogaster* GRs by Kalign [198]with default settings. Poorly-aligned and variable N-terminal and C-terminal regions, as well as several internal regions of highly variable sequences, were excluded from the phylogenetic analysis. Other regions of potentially uncertain alignment between these highly divergent proteins were retained, as removing these regions could potentially compromise subfamily relationships. Based on the trimmed alignment, a PhyML tree was constructed using the substitution model of LG determined by ProtTest [199]. Here, the SH-like local support method was used to assess the significance of phylogenetic clustering.

The *D. tinctorium and L. deliense* iGluRs and IRs were manually annotated according to a TBLASTN (E-value cutoff, <1 × 10^−3^) search against the *D. tinctorium and L. deliense* genome assemblies using all iGluRs and IRs identified by Croset *et al.* [217] across vertebrates and invertebrates, as well as those identified in the recent *T. mercedesae* genome project [98]. Iterative searches were also conducted with termite iGluRs and IRs as queries until no new genes were identified in each major subfamily or lineage. In the phylogenetic analysis, all manually annotated *D. tinctorium and L. deliense* IRs and iGluRs were aligned with the *D. melanogaster* IRs and iGluRs by Mafft [197] using default settings. Phylogenetic analysis proceeded as for the GR genes described above, using the more conserved iGluRs to root the tree.

Reference opsin genes were collected based on the work of Nagata *et al.* [96]. Opsin-like sequences in *M. occidentalis, I. scapularis* and *T. urticae* were obtained from the NCBI database. These opsin genes were classified by phylogenetic analysis using the neighbour-joining method with 1,000 bootstraps. The multiple alignment of the amino acid sequences was carried out using Mafft [197] with the ‘–auto’ option. The gaps deletion of the alignment was set to 75% in MEGA7 [201].

### Prediction of allergenic gene families

Allergenic protein-coding genes in the genomes of acariform mites (*D. tinctorium, L. deliense, T. urticae, D. farinae, S. scabiei*, and *E. maynei*) were predicted using a standalone version of Allerdictor (v. 1.0) [218]. Because the predicted proteome of *D. farinae* is not publicly available, we used protein sequences identified from a new Trinity (v. 2.4) [219] assembly for the prediction of allergenic genes. The allergenic gene clusters were constructed using OrthoMCL (v. 1.4) [207] and individual protein sequences were submitted to Pfam (EBI, v.31.0 [220]) to be assigned to protein families. Mapped Pfam domain identifications were then searched against the AllFam database [112] to retrieve corresponding AllFam identifiers for allergen families. Venn diagrams were constructed using InteractiVenn [221].

### Salivary and cement protein analysis

A total of 159 non-redundant tick cement proteins were retrieved from the NCBI database and then clustered with all *D. tinctorium* and *L. deliense* amino-acid sequences using OrthoMCL in order to identify the tick cement orthologues in the new genomes. Salivary proteins of *I. scapularis* and *T. urticae* have been previously identified using proteomic methods by Kim *et al.* [120] and Jonckheere *et al.* [126], respectively. Proteins that might be present in the saliva of *D. tinctorium* and *L. deliense* were identified by clustering all predicted protein-coding sequences in the new genomes with these *T. urticae* and *I. scapularis* salivary proteins using OrthoMCL [207]. Venn diagrams were constructed using InteractiVenn [221].

### Sample preparation for proteomics

Engorged larvae of *L. deliense* (10 specimens) were pooled from infested rodent specimens (*Bandicota indica, Bandicota savilei* and *Rattus tanezumi*) captured across several provinces in Thailand during the field studies of the CERoPath project [177]. The chigger samples were fixed in 70% ethanol and identified as *L. deliense* using autofluorescence microscopy [222]. For *D. tinctorium*, proteomic analysis was performed on a single ethanol-fixed individual from the same collection used for genome sequencing.

The chiggers were washed with chilled 50 mM ammonium bicarbonate. Soluble protein extracts were prepared by homgenisation in 0.1% w/v, Rapigest (Waters) in 50 mM ammonium bicarbonate using a polypropylene mini-pestle. This was followed by three cycles of sonication on ice (Vibra-cell 130PB sonicator at 20 Hz with microprobe; 10 sec of sonication alternating with 30 sec of incubation on ice). Samples were centrifuged at 13,000 × g for 10 min at 4°C. The supernatant was removed and retained. The *D. tinctorium* individual was homogenised using a mini-pestle in lysis buffer (4% SDS, Tris-hydrochloride, pH 7.6) and sonicated and centrifuged as above. Supernatants from both mite preparations were stored at −80°C.

Protein concentrations of the samples were determined using a Bradford protein assay (Thermo Fisher Scientific). The *L. deliense* protein extract was reduced with 3 mM dithiothreitol (Sigma) at 60°C for 10 min, cooled, then alkylated with 9 mM iodoacetamide (Sigma) at room temperature for 30 min in the dark; all steps were performed with intermittent vortex-mixing. Proteomic-grade trypsin (Sigma) was added at a protein:trypsin ratio of 50:1 and incubated at 37°C overnight. Rapigest was removed by adding trifluoroacetic acid (TFA) to a final concentration of 0.5% (v/v). Peptide samples were centrifuged at 13,000 × g for 30 min to remove precipitated Rapigest. The *D. tinctorium* protein extract was reduced, alkylated and digested with trypsin using the filter-aided sample preparation approach [57]. Peptides from *D. tinctorium* were split into eight fractions using the Pierce High pH Reversed-Phase Peptide Fractionation Kit according to the manufacturer’s instructions. Each digest and fraction was concentrated and desalted using C18 Stage tips (Thermo Fisher Scientific), then dried using a centrifugal vacuum concentrator (Eppendorf) and re-suspended in a 0.1% (v/v) TFA, 3% (v/v) acetonitrile solution.

### Mass spectrometry

Peptides were analysed by on-line nanoflow LC using the Ultimate 3000 nano system (Dionex/Thermo Fisher Scientific). Samples were loaded onto a trap column (Acclaim PepMap 100, 2 cm × 75 μm inner diameter, C18, 3 μm, 100 Å) at 9μl /min with an aqueous solution containing 0.1 %( v/v) TFA and 2% (v/v) acetonitrile. After 3 min, the trap column was set in-line to an analytical column (Easy-Spray PepMap^®^ RSLC 50 cm × 75 μm inner diameter, C18, 2 μm, 100 Å) fused to a silica nano-electrospray emitter (Dionex). The column was operated at a constant temperature of 35°C and the LC system was coupled to a Q-Exactive mass spectrometer (Thermo Fisher Scientific). Chromatography was performed with a buffer system consisting of 0.1 % formic acid (buffer A) and 80 % acetonitrile in 0.1 % formic acid (buffer B). The peptides were separated by a linear gradient of 3.8 – 50 % buffer B over 90 minutes (*D. tinctorium* and *L. deliense* whole digests) or 30 min (*D. tinctorium* fractions) at a flow rate of 300 nl/min. The Q-Exactive was operated in data-dependent mode with survey scans acquired at a resolution of 70,000 at m/z 200. Scan range was 300 to 2000m/z. Up to the top 10 most abundant isotope patterns with charge states +2 to +5 from the survey scan were selected with an isolation window of 2.0 Th and fragmented by higher energy collisional dissociation with normalized collision energies of 30. The maximum ion injection times for the survey scan and the MS/MS scans were 250 and 50 ms, respectively, and the ion target value was set to 1 × 10^6^ for survey scans and 1 × 10^4^ for the MS/MS scans. The MS/MS events were acquired at a resolution of 17,500. Repetitive sequencing of peptides was minimized through dynamic exclusion of the sequenced peptides for 20 sec.

### Protein identification, quantification and enrichment analysis

Thermo RAW files were imported into Progenesis LC–MS (version 4.1, Nonlinear Dynamics). Peaks were picked by the software using default settings and filtered to include only peaks with a charge state between +2 and +7. Spectral data were converted into .mgf files with Progenesis LC–MS and exported for peptide identification using the Mascot (version 2.3.02, Matrix Science) search engine as described above. Tandem MS data were searched against a database including translated ORFs from either the *D. tinctorium* genome (DinoT_V2_aug2016, 19,258 sequences; 8,386,445 residues) and a contaminant database (cRAP, GPMDB, 2012) (119 sequences; 40,423 residues), or the *L. deliense* genome (L_deliense_V2_Aug16, 15,096 sequences; 5,183,596 residues), the *Rattus norvegicus* genome (UniProt, Apr16 7,948 sequences; 4,022,300 residues) and a contaminant database (cRAP, GPMDB, 2012) (119 sequences; 40,423 residues). The search parameters were as follows: the precursor mass tolerance was set to 10 ppm and the fragment mass tolerance was set as 0.05 Da. Two missed tryptic cleavages were permitted. Carbamidomethylation (cysteine) was set as a fixed modification and oxidation (methionine) set as a variable modification. Mascot search results were further validated using the machine learning algorithm Percolator embedded within Mascot. The Mascot decoy database function was utilised and the false discovery rate was <1%, while individual percolator ion scores >13 indicated identity or extensive homology (P <0.05). Mascot search results were imported into Progenesis LC–MS as XML files. Fractions were combined using the Progenesis “combine analysed fractions” workflow. Relative protein abundance was calculated by the Hi-3 default method in Progenesis. Mass spectrometric data were deposited to the ProteomeXchange Consortium (http://proteomecentral.proteomexchange.org) via the PRIDE partner repository [223] with the dataset identifier PXD008346.

Enrichment of protein domains was assessed using Pfam (EBI, v.27.0 [220]) as previously described [224] using the gathering threshold as a cut-off. Briefly, a hypergeometric test for enrichment of Pfam domains in the observed proteome (for identifications supported by ≥2 unique peptides only) relative to the complete search database was performed using R (phyper). The Benjamini & Hochberg step-up false-discovery rate-controlling procedure was applied to the calculated *P* values [225], and enrichment was considered statistically significant where *P* <0.05.

## Availability of data and materials

The data sets supporting the results of this article are available in the GigaDB repository associated with this publication.

**Abbreviations**
CTLD: C-type lectin domain
ERV: endogenous retrovirus
FLVCR: feline leukemia virus subgroup C receptor-related protein
GR: gustatory receptor
IMD: immune deficiency
IR: ionotropic receptor
iGluR: ionotropic glutamate receptors
LGT: lateral gene transfer
MYA: million years ago
NCBI: National Center for Biotechnology Information
OBP: odorant-binding protein
PE: paired-end
PERK: PKR-like endoplasmic reticulum kinase
PGRP: peptidoglycan recognition protein
SMOC: SPARC-related modular calcium-binding protein
TFA: trifluoroacetic acid
XIAP: X-linked inhibitor of apoptosis protein

## Animal ethics

Wild rodents trapped during the CERoPath project [177] and used as a source of chigger material were euthanized by inhaled anaesthetic overdose according to guidelines published by the American Veterinary Medical Association Council on Research [226] and the Canadian Council on Animal Care [227].

## Authors’ contributions

Conceptualisation: A.C.D., B.L.M. Formal analysis: X.D., D.X., S.D.A., Y.F., A.C.D. Funding acquisition: J.W.M., A.C.D., B.L.M. Investigation: K.C., S.D.A., M.J.D. Project administration: A.C.D., B.L.M. Supervision: A.C.D., J.W.M., T.K., B.L.M. Validation: X.D., A.C.D., D.X., B.L.M. Visualization: X.D., D.X., S.D.A., A.C.D., B.L.M. Writing (original draft): B.L.M., X.D. Writing (review and editing): B.L.M., X.D., K.C., T.K., M.J.D., D.X., S.D.A., A.C.D. All authors have read and approved the final manuscript.

## Acknowledgements

The authors are grateful to Serge Morand (Kasetsart University, Thailand) for managing the CERoPath project field collections that provided chigger material and to Joanna Mąkol (Wrocław University of Environmental and Life Sciences, Poland) for identification of the *D. tinctorium* specimens. We also thank the core research staff at the Centre for Genomic Research (University of Liverpool) for DNA library preparations, sequencing and initial data quality control.

## Additional files

**Additional file 1**: Genome assembly and gene set statistics compared with 14 other arachnids (table in .xlsx format).

**Additional file 2**: *K*-mer distributions for *Leptotrombidium deliense* (*A*) and *Dinothrombium tinctorium* (*B*) plotted by GenomeScope (figure in .pdf format).

**Additional file 3**: Identification of repetitive sequences in the *Dinothrombium tinctorium* and *Leptotrombidium deliense* assemblies compared with other acariform mites (table in .pdf format).

**Additional file 4**: The number of gene families shared among acariform mites (*Dinothrombium tinctorium, Leptotrombidium deliense, Tetranychus urticae* and *Sarcoptes scabiei*); alongside other references including *Drosophila melanogaster, Apis mellifera, Tropilaelaps mercedesae, Metaseiulus occidentalis, Ixodes scapularis, Stegodyphus mimosarum* and *Caenorhabditis elegans* by the OrthoMCL classification algorithm (figure in .pdf format).

**Additional file 5**: Gene family contraction and expansion in 11 species of Ecdysozoa (figure in .pdf format).

**Additional file 6**: Changes of gene family size in trombidid mites in comparison with three other acariform mites (table in .xlsx format).

**Additional file 7**: High-confidence protein identifications and abundance scores for *Leptotrombidium deliense* engorged larvae (spreadsheet in .xlsx format).

**Additional file 8**: High-confidence protein identifications and abundance scores for a single adult *D. tinctorium* individual (spreadsheet in .xlsx format).

**Additional file 9**: Phylogeny of carotenoid synthases-cyclases from trombidid mites, spider mites, aphids and fungi (figure in .pdf format).

**Additional file 10**: Genomic scaffold of *Dinothrombium tinctorium* containing a putative lateral gene transfer adjacent to an incontrovertible mite gene (figure in .pdf format).

**Additional file 11**: Microbial reads identified in the trombidid genomic data by the Kraken taxonomic sequence classification system (table in .pdf format).

**Additional file 12**: Peptides detected by mass spectrometry from two terpene synthases in an adult specimen of *Dinothrombium tinctorium* (figure in .pdf format).

**Additional file 13**: Phylogeny of reverse ribonuclease integrases in *Dinothrombium tinctorium* and their closest homologues in other taxa (figure in .pdf format).

**Additional file 14**: Phylogeny of Pol-like polyproteins in trombidid mites and their closest homologues in other taxa (figure in .pdf format).

**Additional file 15**: RNA families identified in the Rfam database in 10 arthropod genomes (table in .xlsx format).

**Additional file 16**: Phylogeny of Dscam protein-coding sequences in *Dinothrombium tinctorium, Leptotrombidium deliense, Tetranychus urticae* and *Ixodes scapularis* (figure in .pdf format).

**Additional file 17**: Phylogeny of peptidoglycan recognition protein sequences in *Dinothrombium tinctorium, Leptotrombidium deliense, Tetranychus urticae* and *Ixodes scapularis* alongside homologous sequences from insects.

**Additional file 18**: Phylogeny of PKR-like endoplasmic reticulum kinase sequences in *Leptotrombidium deliense* and their closest homologues in other taxa (figure in .pdf format).

**Additional file 19**: Predicted allergens in the *Leptotrombidium deliense* genome with top BLAST hit, Pfam domains and AllFam classifications (spreadsheet in .xlsx format).

**Additional file 20**: Orthologous clusters of tick cement proteins in the genomes of *Dinothrombium tinctorium* and *Leptotrombidium deliense* (table in .pdf format).

